# Meta-analysis on reporting practices as a source of heterogeneity in *in vitro* cancer research

**DOI:** 10.1101/2021.10.05.463182

**Authors:** Timo Sander, Joly Ghanawi, Emma Wilson, Sajjad Muhammad, Malcolm Macleod, Ulf Dietrich Kahlert

**Author notes:** Corresponding author mail address.

## Abstract

**Background:** Heterogeneity of results of exact same research experiments oppose a significant socio-economic burden. *In vitro* research presents the early step of basic science and drug development projects. Insufficient methodological reporting is likely to be one of the contributors to results heterogeneity, however, little knowledge on reporting habits of *in vitro* cancer research and their effects on results reproducibility is available. Glioblastoma is a form of brain cancer with largely unmet clinical need.

**Methods:** Here we use systematic review to describe reporting practices in *in vitro* glioblastoma research using the U87-MG cell line and perform multilevel random-effects meta-analysis followed by meta-regression to explore sources of heterogeneity within that literature, and any associations between reporting characteristics and reported findings.

**Results:** In 137 identified articles, the overall methodological reporting is disappointing, e.g., the control type, mediums glucose level and cell density are reported in only 36.5, 21.2 and 16.8 percent of the articles, respectively. After adjustments for different drug concentrations and treatment durations, a three-level meta-analysis proves meaningful results heterogeneity across the studies (*I*^*2*^ = 70.1%).

**Conclusions:** Our results further support the ongoing efforts of establishing consensus reporting practices to elevate durability of results. By doing so, we hope that this work will raise awareness of how stricter reporting may help to improve the frequency of successful translation of preclinical results into human application, not only in neuro-oncology.

**Funding:** We received no specific funding for this project.

## Introduction

Progress in scientific research is a dynamic process which thrives in the interaction of diverse research groups addressing shared problems. The scientific model has new findings either confirmed or refuted by other scientists, so that science becomes self-correcting (Merton, 1973). However, one key foundation for self-correction is that key experimental methods needed for interpretation and repetition of published research are described in sufficient detail. Recent efforts to replicate key findings in cancer biology and further research areas have raised scientific, ethical and economic concerns (Begley & Ellis, 2012; Begley C. Glenn & Ioannidis John P.A., 2015; Freedman et al., 2017; Global Biological Standards Institute (GBSI), 2013; Hirsch & Schildknecht, 2019; Jarvis & Williams, 2016; Prinz et al., 2011; Wen et al., 2018).

Glioblastoma is a malignant brain tumour with a median time of survival of around 15 months (Tamimi & Juweid, 2017). First-line treatment consists of a multimodal approach of surgical resection followed by radiation therapy and chemotherapy with the alkylating agent temozolomide (TMZ) (Tan et al., 2020). A previous systematic review of the *in vivo* literature describing the efficacy of TMZ showed limited reporting of key study design features and low prevalence of reporting of measures to reduce risks of bias (Hirst et al., 2013). *In vitro* glioblastoma research commonly uses the commercially available cell line *Uppsala-87 Malignant Glioma* (U-87 MG) (Robertson et al., 2019; Poon et al., 2021), originally derived in 1966 from a 44-year-old female patient at Uppsala University (Pontén & Macintyre, 1968). However, the currently available U-87 MG line distributed by the American Type Culture Collection (ATCC, Manassas, Virginia) (HTB-14™, ATCC®, 2021) has been found to be different to the original version (Allen et al., 2016). It is unclear to what extent these U-87 MG cells are truly representative of the original tumour tissue and whether they allow for reproducible experiments when serving as glioblastoma models.

The reproducibility of *in vitro* glioma research is likely to depend on the completeness of reporting of key study design features including reporting of risks of bias. Here we use a systematic review to portray the *in vitro* literature describing the effectiveness of TMZ in reducing the growth of U-87 MG cells, with a focus on the use of clinically relevant drug concentrations and treatment durations, on methodological reporting and how this might influence the reproducibility of results.

## Methods

The study protocol is available at the Open Science Framework (https://osf.io/9k3dq) and was uploaded before full text based screening and data extraction began. Deviations from the protocol are described in the methods section.

### Systematic Review

#### Systematic literature search and screening

The systematic search was conducted on the databases PubMed, Embase and Web of Science on the 26^th^ of August 2020 using the search strategy described in Supplement 1. Two reviewers independently screened article titles and abstracts for potential inclusion, with discrepancies resolved by a third reviewer. This was followed by full text screening. We included studies describing controlled *in vitro* cell culture experiments that compared the effect of a single TMZ treatment on the viability of U-87 MG cells with that in untreated controls. We also required that cell viability was measured by colorimetric assay or by cell counting; and that the authors used Dulbecco’s Modified Eagle Medium (DMEM) as the ATCC recommends using a modified Eagle medium for U-87 MG cell cultures (HTB-14™, ATCC®, 2021). We only included original peer-reviewed research articles in the English language, with no restrictions made on publication year. Our protocol had included consideration of cell growth rates in xenotransplantation models, but we later decided to focus exclusively on *in vitro* research. Inclusion and exclusion criteria are given in Supplement 2.

#### Data extraction

We recorded effect sizes for change in cell viability in response to TMZ compared to untreated control, TMZ concentration and duration of exposure. We recorded fifteen experimental parameters (Supplement 3); eight risks of bias items (Supplement 4); and the journal impact factor (JIF) for that journal in the year of publication (Web of Science Group, 2020). Consideration of JIF had not been included in our study protocol and should be considered an exploratory analysis.

Two reviewers independently recorded study design features, risk of bias items and effect sizes, with reconciliation of discrepancies by a third reviewer where necessary; data for effect sizes were reconciled if they differed by more than 15% of the largest effect size; otherwise, the mean of the extracted values from the two reviewers was used. Where information was missing, we contacted authors for clarification; to reduce the burden on them to reply we limited this request to a maximum of eleven items (Supplement 5).

#### Reporting quality of experimental parameters and articles

Where an article reported multiple relevant experiments, a parameter was considered as reported for an article if it was reported in every belonging experiment. When calculating the number of items reported for each article, we did not consider the volume of added TMZ and of control fluid as we considered this was included if the drug and control concentration was given. We analysed change in reporting quality over time, and any relationship between reporting quality and JIF, using linear regression.

### Meta-analysis

#### Exclusions from the meta-analysis

We excluded experiments that did not report data essential for our analysis such as cell viability data for the untreated control group, TMZ concentration, duration of treatment or the number of experimental units; and we excluded baseline data (where treatment duration = 0).

#### Effect size

We calculated a cell viability reduction caused by TMZ compared to the corresponding untreated control as raw mean difference with all data given in relation to the control viability as

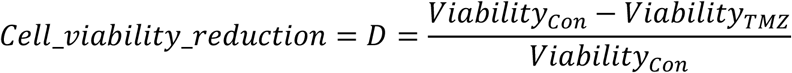

with its variance

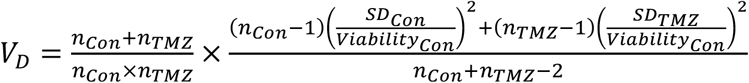

where n_Con_ and n_TMZ_ represented the number of experiments in the control and TMZ group, respectively, and SD_Con_ and SD_TMZ_ represented the standard deviation (SD) of cell viabilities in the control and TMZ group. If the variance in the control group was not reported, we assumed this to be equivalent to variance in the corresponding TMZ group. If it was not clear whether variance was reported as SD or standard error of the mean (SEM), we assumed they were SEM, as a more conservative approach.

#### Multi-level random-effects meta-analysis

We used random-effects meta-analysis (Riley et al., 2011). Because a single article might contribute several effects sizes, we used a three-level model where the first level represented the raw cell viability data, the second level all the effects from a given article and the third level the article itself. This accounts for the relative non-independence of effects reported in the same article (Cheung, 2013). Moreover, as the exact correlations of the dependent effects within an article were unknown, we used robust-variance-estimation (Hedges et al., 2010).

To estimate tau^2^, we used the restricted-maximum-likelihood method (Harville, 1977). This method has recently been shown to be robust for non-normally distributed effect sizes (Langan et al., 2019) and is recommended for the estimation of tau^2^ (Viechtbauer, 2005; Langan et al., 2019). We used the t-distribution for the calculation of the weighted mean effect as this accounts for uncertainty in the estimation of tau^2^ (Higgins et al., 2009). We took the within-level-three estimate of tau^2^ as a measure of reproducibility of findings between studies and subsequently as an indicator of irreproducibility of results across articles.

#### Meta-regression

We tested ten parameters in univariable meta-regression which were defined a priori. For these, a reduction of within-level-three tau^2^ would indicate that that parameter moderated reproducibility, with lower within-level-three tau^2^ indicating that findings would be more likely to reproduce if that parameter was controlled between the original and replicating experiment. We took the same approach to establish any effect of articles overall reporting quality or JIF. We also used univariable meta-regression to analyse the effects of TMZ dose and treatment duration. We transformed TMZ concentrations into a four-parameter log-logistic dose-response model (Hill, 1910). A similar four-parameter log-logistic time-response model was built for the treatment durations. Finally, we conducted multivariable meta-regression of the effect of dose, duration and moderators proven significant in univariable meta-regression. In all analysis we set a significance level of 0.05

#### Software

To remove duplicate articles from the systematic search results we used two approaches, the deduplication function integrated in Zotero (Roy Rosenzweig Center for History and New Media, 2021) and one developed by the CAMARADES group (Hair, 2019). Articles were removed if they were detected as a duplicate by both functions. Afterwards, articles were manually screened for remaining duplicates. Screening and data extraction used the Systematic Review Facility (SyRF) for preclinical systematic reviews (Bahor et al., 2021). Graphically presented data were extracted with the WebPlotDigitizer (Ankit Rohatgi, 2020). Meta-analysis was performed within the RStudio environment (RStudio Team, 2020) using the programming language R (R Core Team, 2020). We used the rma.mv function of the R package Metafor (Viechtbauer, 2010) for multi-level meta-regressions, the R package clubSandwich (Pustejovsky, 2021) for robust-variance-estimation, the R package orchard (Nakagawa et al., 2021) for the calculation of marginal *R*^*2*^ and *I*^*2*^, the drm function of the R package drc (Ritz et al., 2015) for the dose- and time-response models and the lm function of the integrated R package stats (R Core Team, 2020) for linear regressions. The full R code and datasets are available on GitHub (https://github.com/TimoSander/Reporting-practices-as-a-source-of-heterogeneity-in-in-vitro-cancer-research/).

## Results

### Systematic search

We identified 1158 publications of which 137 articles met our inclusion criteria and were included in the systematic review; 101 provided sufficient data to be included in the meta-analysis (Figure 1). The Supplement 6 contains a list of all included articles. These 137 articles described 828 experiments where every different combination of drug concentration and treatment duration used was considered as an individual experiment. The main reason for exclusion from meta-analysis was an unreported number of contributing experimental units (n = 24 articles).

**Figure 1:**
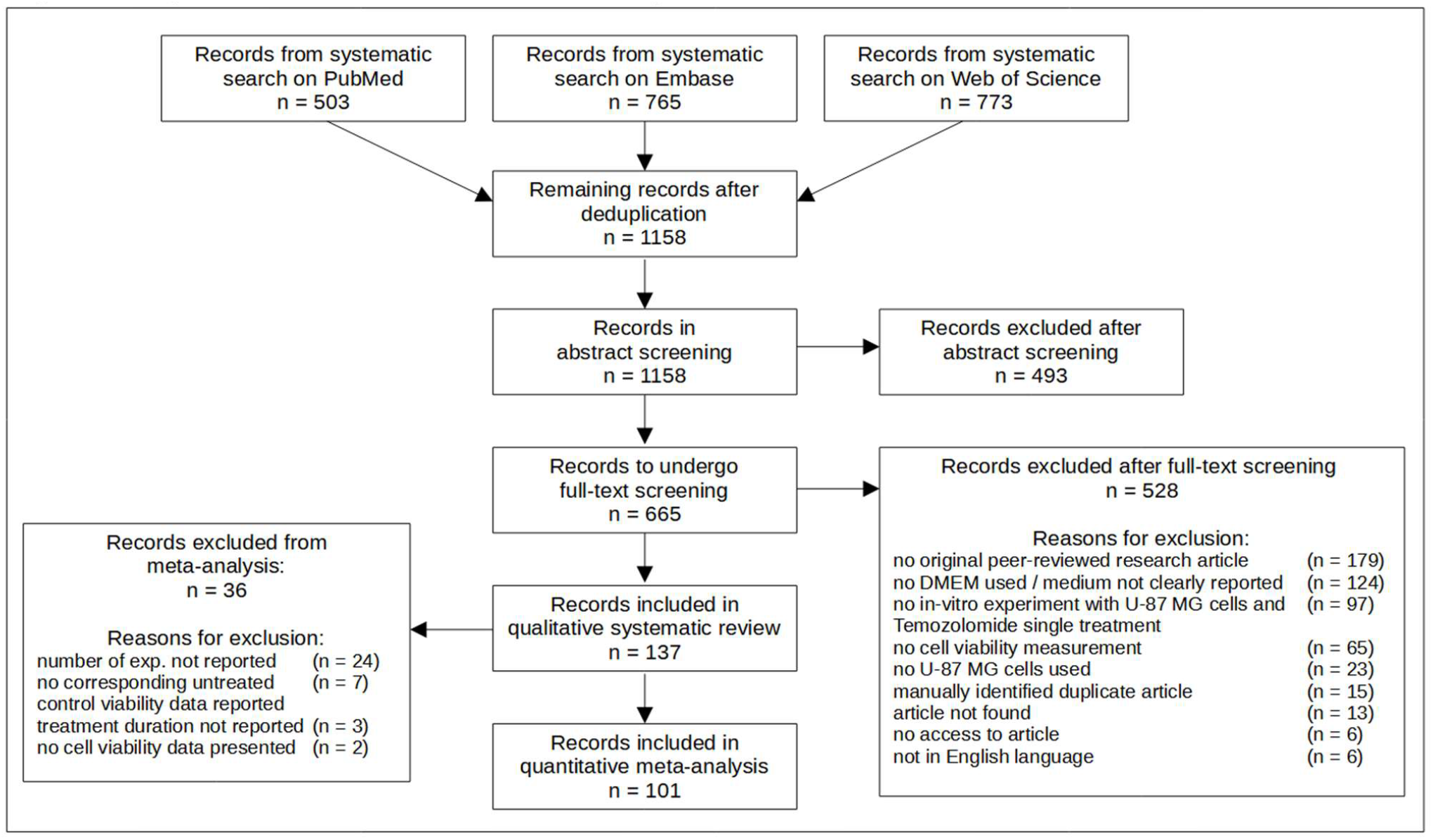
Systematic search and screening results. Presentation based on the PRISMA statement (Page et al., 2021). Systematic searches were conducted in August 2020. Qualitative analysis included all calculations in this paper except meta-analysis and meta-regressions. One reason for exclusion per excluded article. Exp. = experiments; DMEM = Dulbecco’s modified Eagle’s medium; U-87 MG = Uppsala-87 Malignant Glioma.

### Experimental parameters distribution

Across 137 publications, a broad range of experimental characteristics were included. The most common source of U-87 MG cells was the ATCC (66 articles, 48.2% of all included articles; Table 1a). A cell line authentication report was available in 16 articles (11.7%) and eight articles (5.8%) described testing for mycoplasma contamination. The reported cell passage number ranged from three to one hundred (median of 15), but 123 publications (89.8%) did not report it. Only 29 of 137 articles (21.2%) reported the level of culture mediums glucose, and in these high glucose supplementation (4500 mg/dl) was most prevalent (in 24 of 29). Control treatment was dimethyl sulfoxide in 37 and culture medium alone in 13 articles; in 87 it was not reported or only labelled as “untreated control” without further specification. The most common cell viability assessment method was the 3-(4,5-dimethylthiazol-2-yl)-2,5-diphenyltetrazolium (MTT) assay (67 articles, 48.9%). The concentration of U-87 MG cells used ranged from 5 to 500 cells per µl (Table 1b), with a median of 30 cells per µl. Ninety-three articles included information for the number of cells per well but not the volume in which these cells were plated. Cells were passaged based on confluence (13 articles, range from 50 to 90% confluence) or on time (four articles, range from 2 to 7 days), but criteria for cell passaging were not stated in 120 studies (87.6%; Table 1c).

**Tab. 1:**
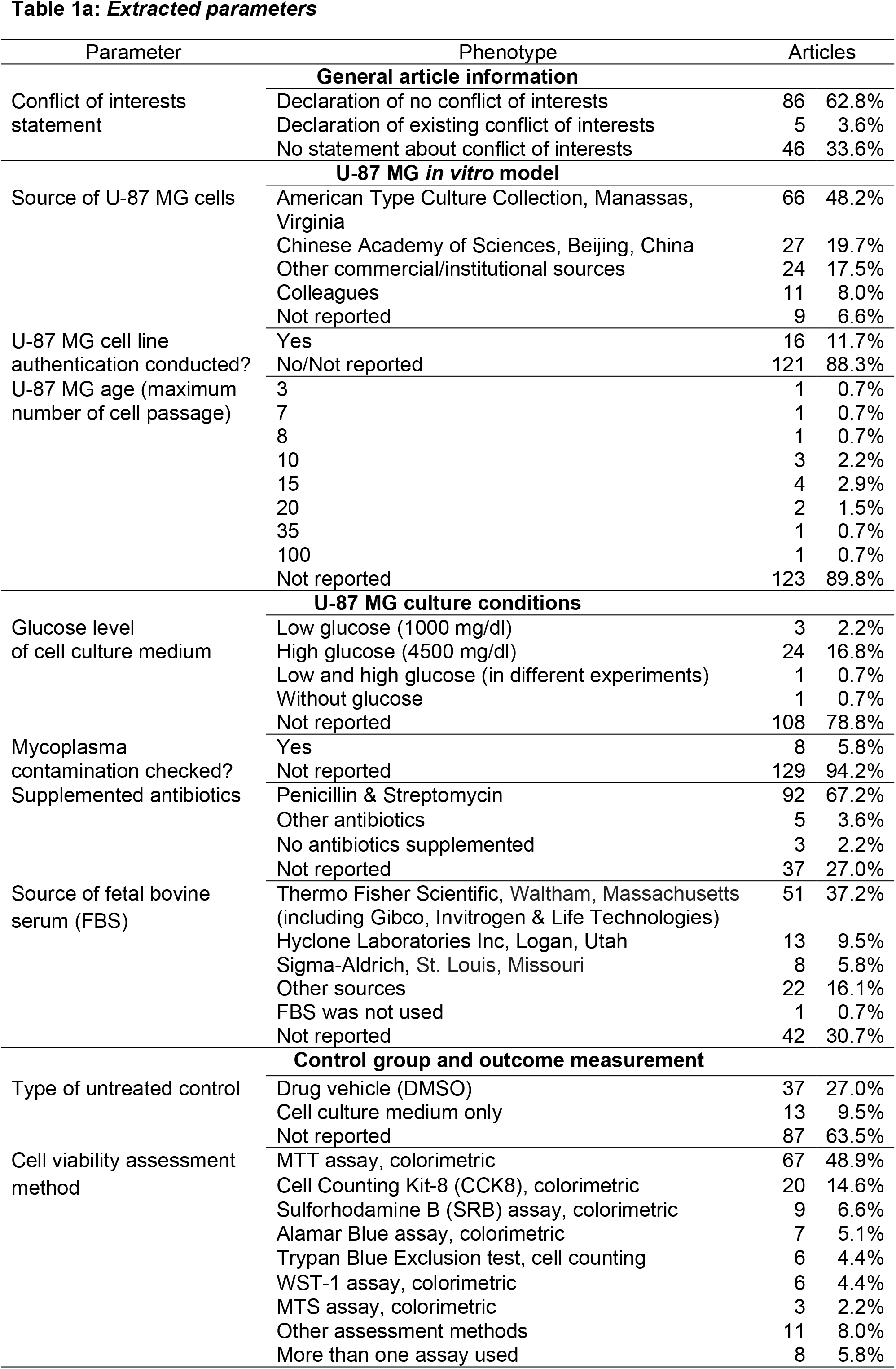
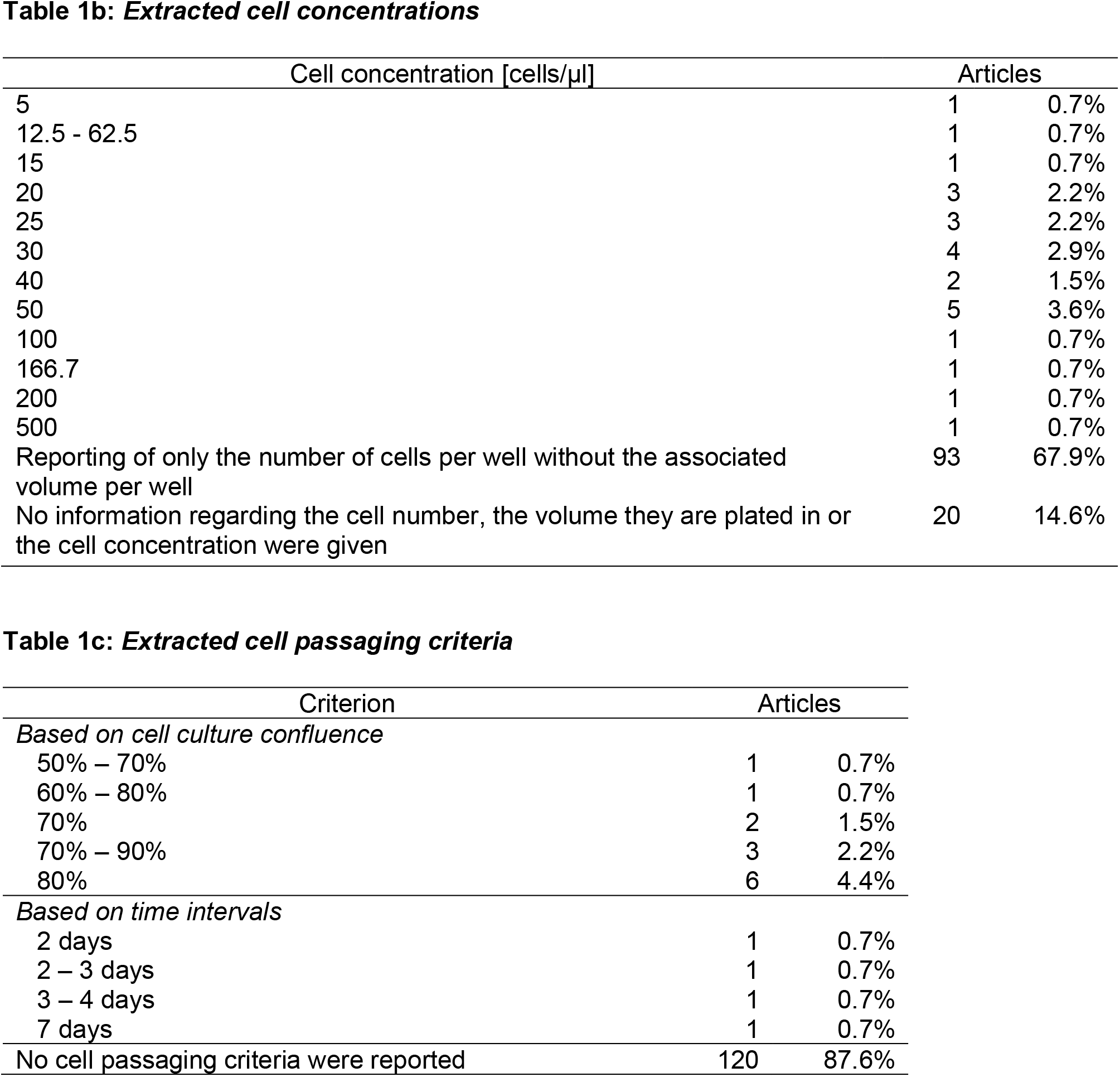
Extracted parameter phenotypes (including additional information obtained through contacting the authors). One phenotype per article. The column “articles” shows the absolute and relative frequencies of articles with the parameter phenotype in relation to all 137 included articles. DMEM = Dulbecco’s modified Eagle’s medium; U-87 MG = Uppsala-87 Malignant Glioma; MTT = 3- (4,5-dimethylthiazol-2-yl)-2,5-diphenyl-2H-tetrazolium bromide; WST-1 = Water-Soluble-Tetrazolium-1; MTS = 3-(4,5-dimethylthiazol-2-yl)-5-(3-carboxymethoxyphenyl)-2-(4-sulfophenyl)-2H-tetrazolium.

Overall, 98 different TMZ concentrations (10nM to 16.0mM, median 100 µM; Supplement 7) and 20 different treatment durations (4 hours to 12 days, median 3 days; Supplement 8) were reported. In several articles it was not clear whether cell viability was measured directly after TMZ exposure, or whether there was a “wash out” or recovery period. In others, it was unclear whether TMZ was added to the cells once and remained in suspension or whether TMZ was added repeatedly at different times. For the purposes of the meta-analysis, we assumed a single TMZ addition with continuous incubation for the reported time followed directly by the assessment of cell viability.

### Completeness of Reporting

Several key experimental parameters were reported in fewer than half of the articles - the type of untreated control was reported in 36.5%, the culture mediums glucose level in 21.2% and U-87 MG cell age in 7.3% of all 137 articles (Figure 2a). The median number of quality items reported was 8.4 of 16 (range from three to thirteen). Analysis of change over time suggested some improvement (0.635% per year, *p* = .011, Figure 2b); and reporting quality seemed to be higher for articles published in journals with higher impact factors (1.74% per unit increase in JIF unit, *p* < .001, Figure 2c).

**Figure 2:**
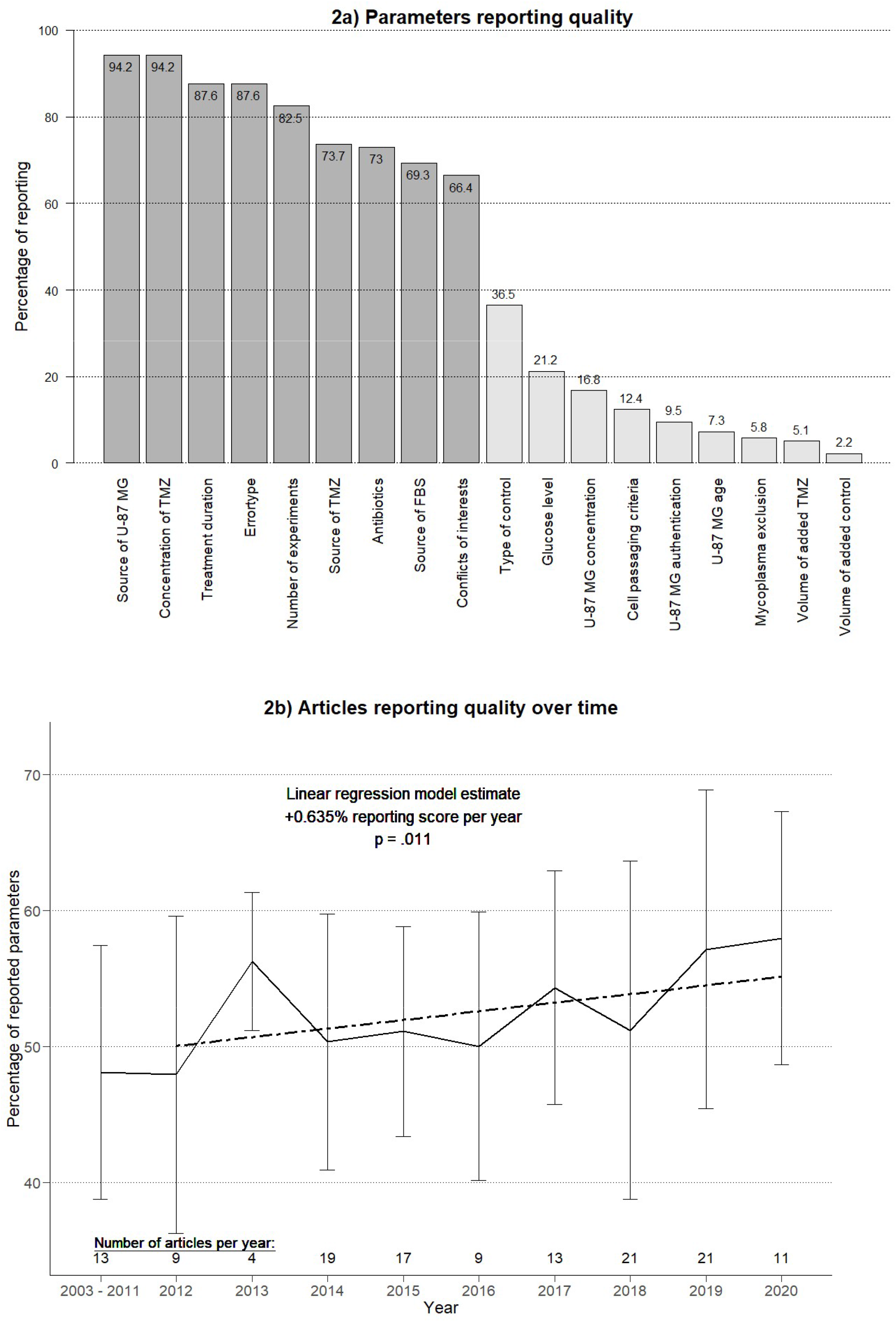
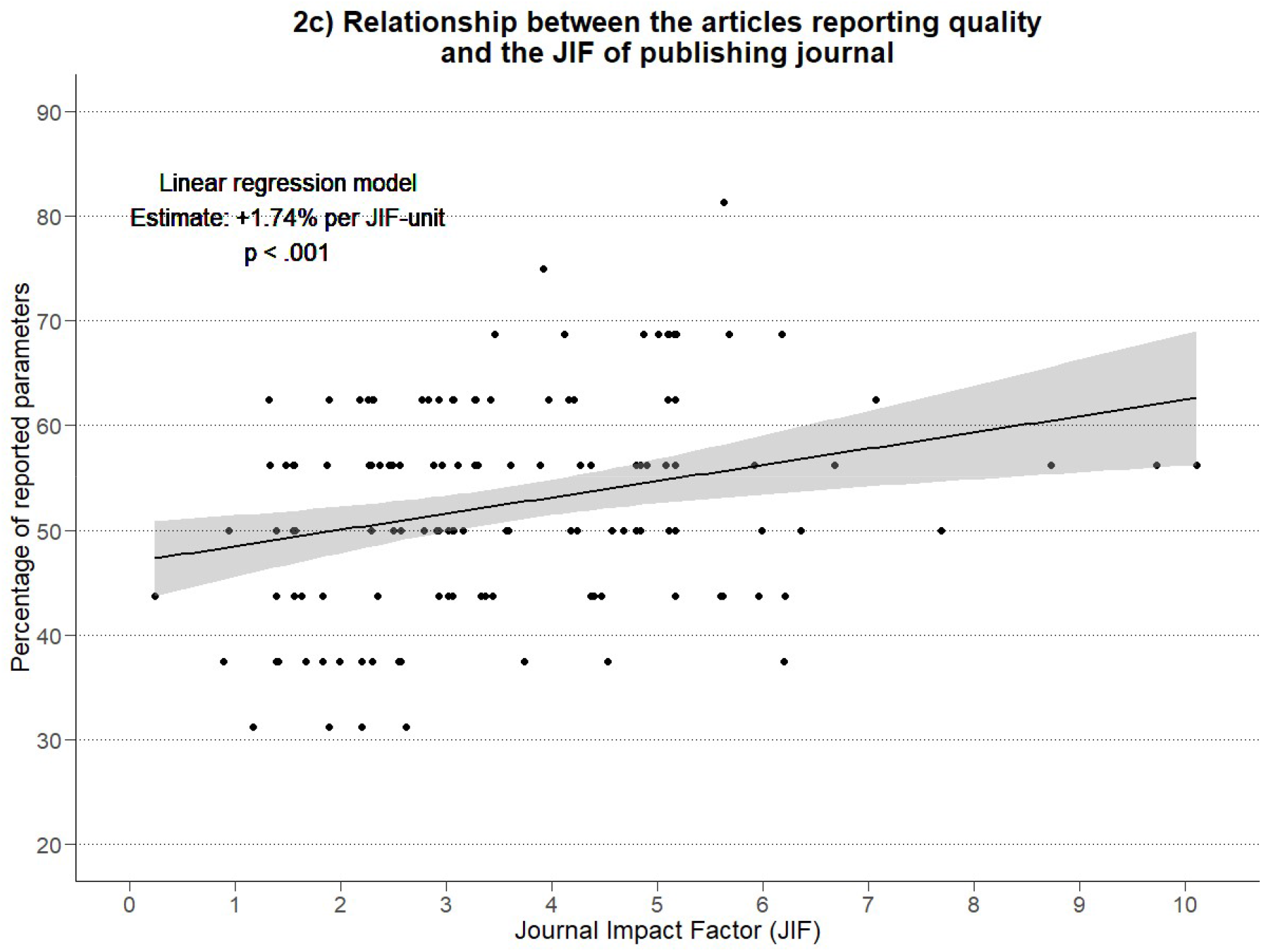
Reporting quality (of parameters, over time and depending on the JIF) **a** The reporting quality of a parameter was defined as the share of articles that reported this parameter phenotype in comparison to all 137 included articles. This share is shown on the top of each bar for each parameter. A parameter was considered to be reported if its phenotype was clear based on the information provided in the original full-text research article. **b** Linear regression model of articles reporting quality (proportion of reported parameters in 16 selected parameters) and year of publication. The articles published before 2012 were summarized because of low numbers of articles published in these years (but the exact years were used for the regression). Only articles until the time of systematic search in August 2020 were considered. The dotted line represents the linear regression line; error bars indicate the standard deviation around the mean reporting score per year represented by the continuous line. **c** Linear regression model of articles reporting quality and the JIF of the articles publishing journal in the year of publication. The grey area marks the 95 % confidence interval of the regression model prediction. JIF were obtained from the Clarivates InCites Journal Citation Reports (Web of Science Group, 2020). For the articles published in 2020, the JIF of 2019 replaced the JIF of 2020 as the more recent was not available at time of analysis. One article was omitted as no JIF could be obtained. U-87 MG = Uppsala 87 Malignant Glioma; FBS = Fetal bovine serum; JIF = Journal impact factor; TMZ = Temozolomide.

### Reporting of measures to reduce risks of bias

Not one of 137 articles described a sample size calculation, random allocation to experimental group, blinded outcome assessment or the use of a pre-registered protocol specifying the hypotheses and outcomes (Table 2). The methods used to calculate cell viability average and error values were unclear in 92 articles, and the number of independent experiments and technical replicates per experiment conducted was unclear in 47 articles. The mean number of measures to reduce risks of bias reported was 2.9.

**Table 2:**
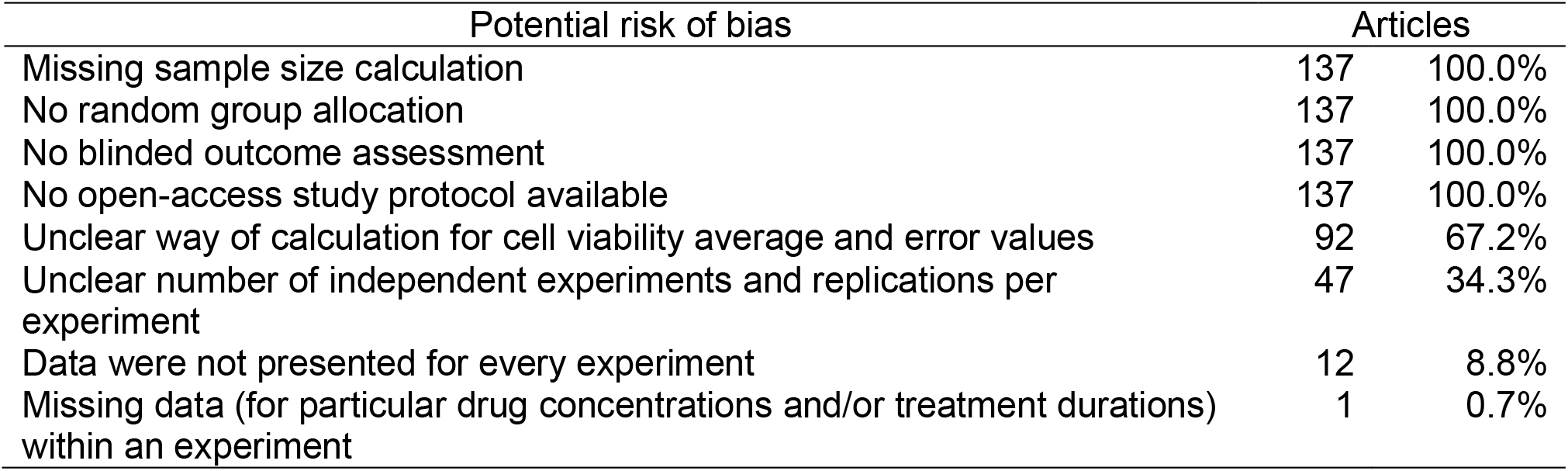
Risk of bias factors prevalence. The column “articles” shows the absolute and relative prevalence of articles having a particular risk of bias factor in comparison to all 137 included articles.

| Potential risk of bias | Articles |  |
| --- | --- | --- |
| Missing sample size calculation | 137 | 100.0% |
| No random group allocation | 137 | 100.0% |
| No blinded outcome assessment | 137 | 100.0% |
| No open-access study protocol available | 137 | 100.0% |
| Unclear way of calculation for cell viability average and error values | 92 | 67.2% |
| Unclear number of independent experiments and replications per experiment | 47 | 34.3% |
| Data were not presented for every experiment | 12 | 8.8% |
| Missing data (for particular drug concentrations and/or treatment durations) within an experiment | 1 | 0.7% |

### Meta-analysis of the effect of TMZ

The observed effect of TMZ is highly heterogeneous; variation within different experiments in the same article (represented by *I*^*2*^ of level-two variance) accounts for 56.6% of observed variance, variation of the effect across different articles (represented by *I*^*2*^ of level-three variance) accounts for 42.9% of observed variance, and the variance due to random chance expected if all experiments of all articles were held under identical conditions made up only 0.5% of the total variance (Table 3). The heterogeneity of results across the articles is reflected in a SD of +/- 16.6% (95% CI for this SD estimate from 13.9% to 19.8%) around a global estimate of a reduction in cell viability following TMZ treatment of 33.8% (95% CI from 30.0 to 37.7%) compared to the untreated control.

**Table 3:** Random-effects three-level meta-analysis suggests significant irreproducibility. Random-effects three-level meta-analysis using the raw data the effects were calculated with as first level, the effect sizes within each article as second level and the articles the effects were reported in as third level. tau^2^: estimator of the variance of true effects (level-two variance = within-articles-variance; level-three variance = between-articles-variance = representant of irreproducibility); tau = square root of tau^2^; *I*^*2*^: proportion of within- and between-articles-variance, respectively, of the total observed variance including sampling error. tau^2^ estimator: restricted-maximum likelihood. Cochran’s Q was used as the test for heterogeneity using a chi-squared distribution; values in square brackets show confidence intervals with significance level set at 0.05. df = degrees of freedom; TMZ = Temozolomide; U-87 MG = Uppsala-87 Malignant Glioma.

| Included data |  |
| --- | --- |
| Number of included effects of TMZ on U-87 MG viability | 644 |
| Number of included articles the effects were reported in | 101 |
| Effect of TMZ |  |
| Overall weighted mean effect of TMZ<br>(U-87 MG viability reduction compared to an untreated control) | 33.8% [30.0%, 37.7%] |
| Investigation of heterogeneity |  |
| Test whether heterogeneity is present |  |
| Q (df = 643): | 134066.1 |
| p value | < .001 |
| Total I<sup>2</sup>: | 99.5% |
| Within-articles-variance of the true effect |  |
| tau<sup>2</sup> | 3.6% [3.2%,4.1%] |
| tau | 19.0% [17.9%,20.3%] |
| I<sup>2</sup> (= proportion of total variance) | 56.6% |
| Between-articles-variance of the true effect (~Irreproducibility) |  |
| tau<sup>2</sup> | 2.8% [1.9%,3.9%] |
| tau | 16.6% [13.9%,19.8%] |
| I<sup>2</sup> (= proportion of total variance) | 42.9% |

### Drivers of heterogeneity

The within-articles-variance of effects reported in the same article could be, as expected, partly explained by differences in TMZ concentrations (50.7%) and treatment durations (5.0%) (Table 4). However, both features did not explain parts of between-articles-variance of effects (Table 5). Combining TMZ concentration and treatment duration in a multivariable meta-regression reduced within-articles-variance tau^2^ from 3.6% to 1.6% while the estimated between-articles-variance increased from 2.8% to 3.8% (Table 6).

**Table 4:**
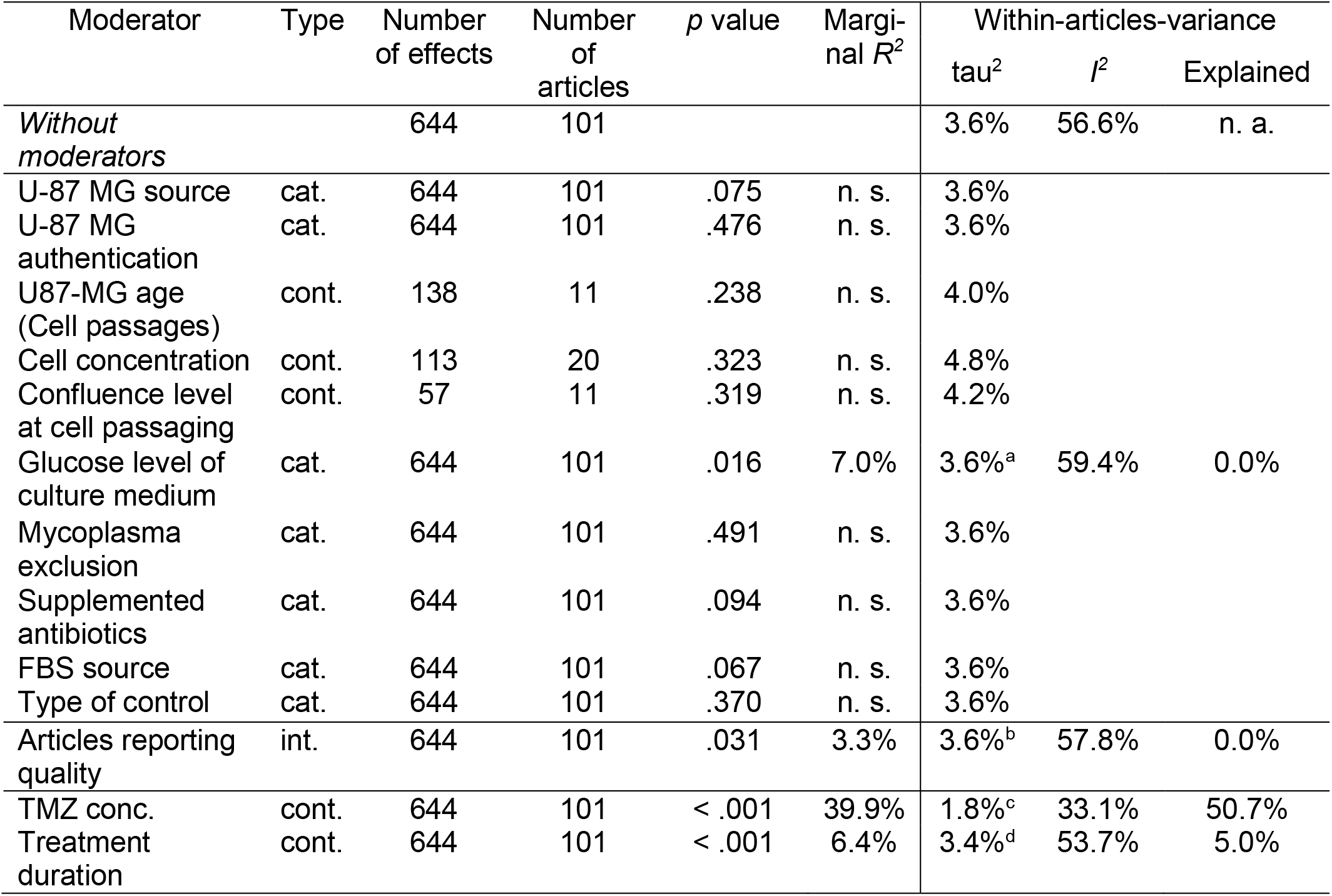
Moderators of within-articles-variance of true effects. Random-effects three-level meta-regressions with the raw data the effects were calculated with as first level, the reported effects as second level and the articles the effects were reported in as third level. Marginal *R*^*2*^ indicates the regression model fit (Nakagawa & Schielzeth, 2013); tau^2^: estimator of the variance of true effects; tau = square root of tau^2^; *I*^*2*^: proportion of within-articles-variance of the total observed variance including sampling error. tau^2^ estimator: restricted-maximum likelihood. The column “explained” indicates the reduction of tau^2^ after including the particular moderator compared to tau^2^ without moderators (only applicable if the number of included effects and articles is identical). Types of moderators: cat. = categorical; cont. = continuous, int. = interval. For some continuous moderators, the number of effects and articles included in the regression is reduced due to non-reporting which leads to a limited comparability of tau^2^ between parameters with different numbers of belonging articles and effects. “Not reported” was included as a category for categorical moderators. The *p* value is for the test of the moderator. *R*^*2*^, *I*^*2*^ and the explained heterogeneity were only calculated for moderators that prove significance in the test of the moderator (alpha = .05). TMZ conc. = Temozolomide concentration. **a**: 95%-confidence-interval (CI): tau^2^: [3.2%,4.1%]; **b**: 95%-CI of tau^2^: [3.2%,4.1%]; **c**: 95%-CI of tau^2^: [1.6%.2.1%]; **d**: 95%-CI of tau^2^: [3.0%,3.9%].

**Table 5:**
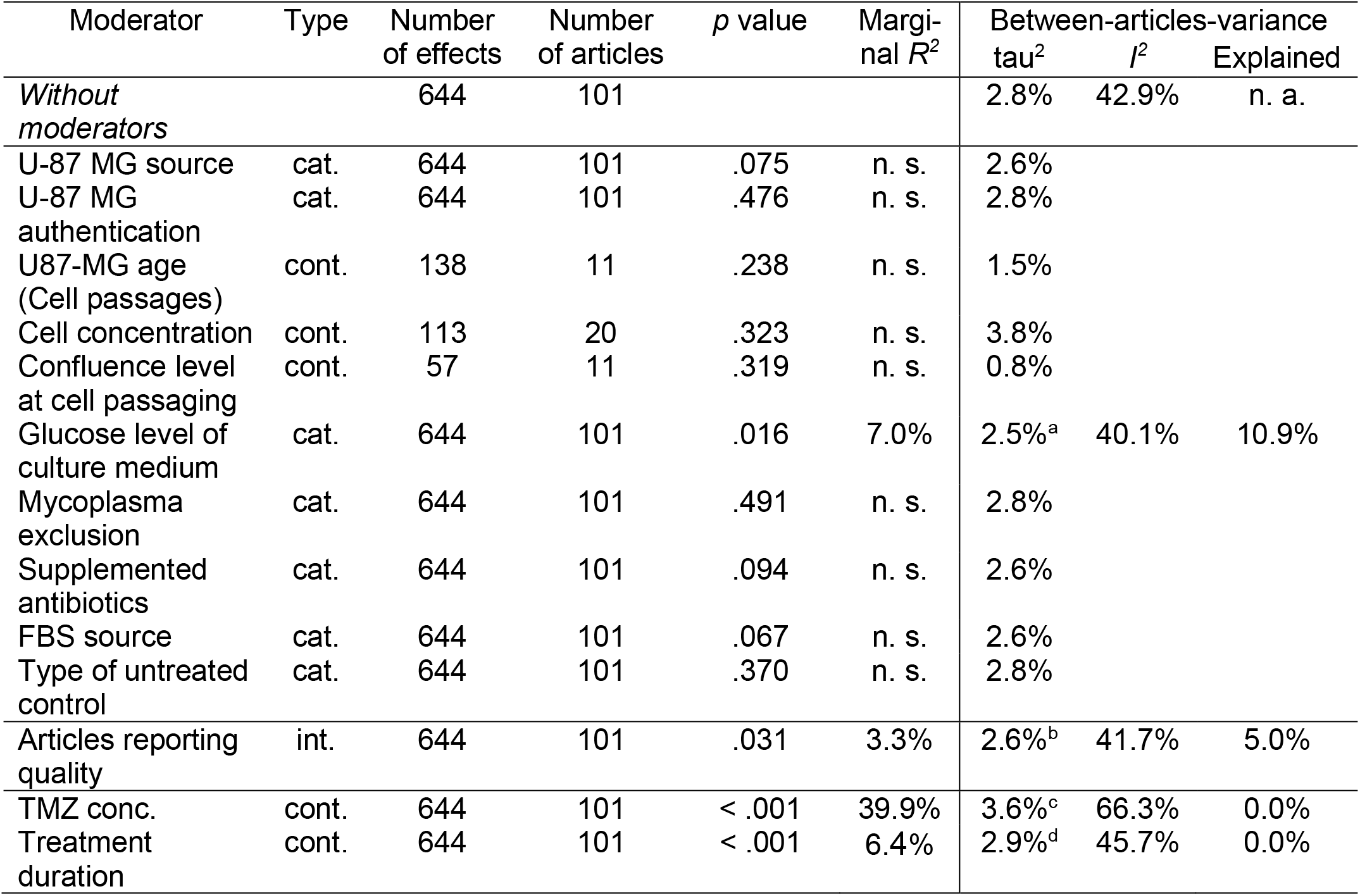
Moderators of between-articles-variance of true effects. Random-effects three-level meta-regressions with the raw data the effects were calculated with as first level, the reported effects as second level and the articles the effects were reported in as third level. Marginal *R*^*2*^ indicates the regression model fit (Nakagawa & Schielzeth, 2013); tau^2^: estimator of the variance of true effects; tau = square root of tau^2^; *I*^*2*^: proportion of between-articles-variance of the total observed variance including sampling error. tau^2^ estimator: restricted-maximum likelihood. The column “explained” indicates the reduction of tau^2^ after including the particular moderator compared to tau^2^ without moderators (only applicable if the number of included effects and articles is identical). Types of moderators: cat. = categorical; cont. = continuous, int. = interval. For some continuous moderators, the number of effects and articles included in the regression is reduced due to non-reporting which leads to a limited comparability of tau^2^ between parameters with different numbers of belonging articles and effects. “Not reported” was included as a category for categorical moderators. The *p* value is for the test of the moderator. *R*^*2*^ and *I*^*2*^ and the explained heterogeneity were only calculated for moderators that prove significance in the test of the moderator (alpha = .05). n. s. = not significant. TMZ conc. = Temozolomide concentration. **a**: 95%-confidence-interval (CI): tau^2^: [1.7%,3.6%]; **b**: 95%-CI of tau^2^: [1.8%,3.8%]; **c**: 95%-CI of tau^2^: [2.6%,5.0%]; **d**: 95%-CI of tau^2^: [2.1%,4.2%]

**Table 6:**
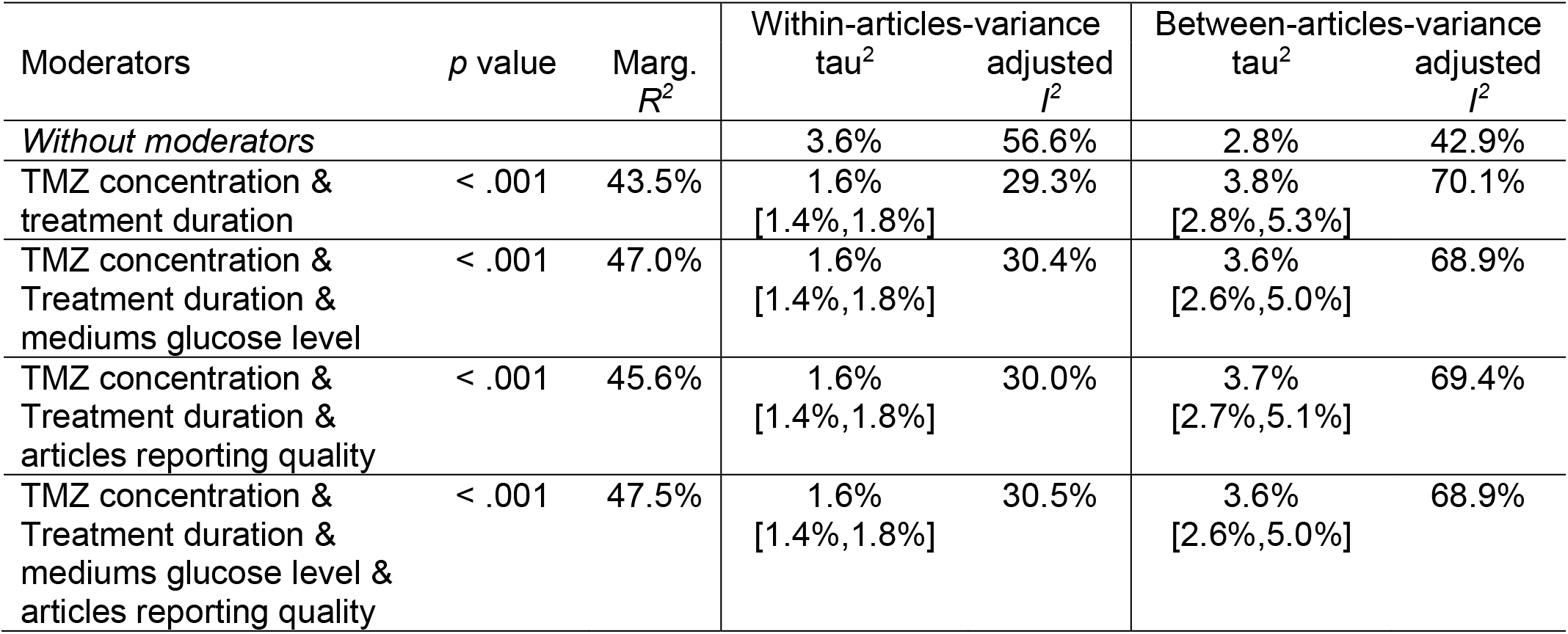
Multivariable meta-regressions. Multivariable random-effects three-level meta-regressions with the raw data the effects were calculated with as first level, the reported effects as second level and the articles the effects were reported in as third level. The *p* value is for the test of the moderator. Marginal *R*^*2*^ indicates the regression model fit (Nakagawa & Schielzeth, 2013); tau^2^: estimator of the variance of true effects; adjusted *I*^*2*^: proportion of within- and between-articles-variance of true effects, respectively, of the total observed variance including sampling error with the indicated moderators. tau^2^ estimator: restricted-maximum likelihood. For all multivariable meta-regressions, 644 effects in 101 articles were included. TMZ = Temozolomide.

| Moderators | *p* value | Marg.<br>*R*<sup>2</sup> | Within-articles-variance<br>tau<sup>2</sup> | adjusted<br>*I*<sup>2</sup> | Between-articles-variance<br>tau<sup>2</sup> | adjusted<br>*I*<sup>2</sup> |
| --- | --- | --- | --- | --- | --- | --- |
| *Without moderators* |  |  | 3.6% | 56.6% | 2.8% | 42.9% |
| TMZ concentration &<br>treatment duration | < .001 | 43.5% | 1.6%<br>[1.4%,1.8%] | 29.3% | 3.8%<br>[2.8%,5.3%] | 70.1% |
| TMZ concentration &<br>Treatment duration &<br>mediums glucose level | < .001 | 47.0% | 1.6%<br>[1.4%,1.8%] | 30.4% | 3.6%<br>[2.6%,5.0%] | 68.9% |
| TMZ concentration &<br>Treatment duration &<br>articles reporting quality | < .001 | 45.6% | 1.6%<br>[1.4%,1.8%] | 30.0% | 3.7%<br>[2.7%,5.1%] | 69.4% |
| TMZ concentration &<br>Treatment duration &<br>mediums glucose level &<br>articles reporting quality | < .001 | 47.5% | 1.6%<br>[1.4%,1.8%] | 30.5% | 3.6%<br>[2.6%,5.0%] | 68.9% |

The glucose level applied in the cell culture medium was the only experimental parameter significantly associated with heterogeneity of results across the articles (*p* = .016, Table 5). The moderator fit indicated by marginal *R*^*2*^ was 7.0%, and 10.9% of the between-articles-variance tau^2^ could be explained because of differences in the glucose level. In other words, roughly 11 percent of heterogeneous results were attributed to different glucose levels used in the culture medium. The different glucose levels and effects are shown in Table 7, indicating a significantly smaller effect of TMZ in the articles using high glucose supplementation (cell viability reduction of 23.1% compared to the untreated control, 95% CI from 15.8 to 30.5%) than in the articles using an unreported glucose level (37.1%, 95% CI from 28.6 to 45.6%, *p* = .002).

**Table 7:**
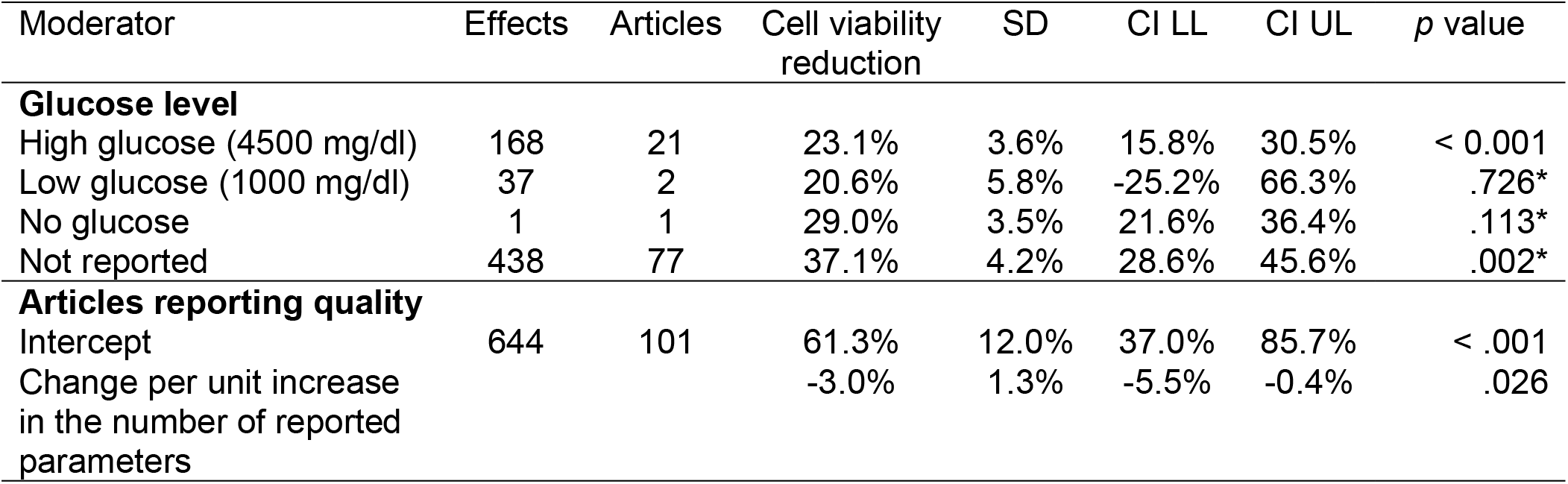
Correlation between the effect and culture mediums glucose level as well as articles reporting quality. Univariable Random-effects three-level meta-regressions with the raw data the effects were calculated with as first level, the reported effects as second level and the articles the effects were reported in as third level. Effects were estimated using robust-variance-estimation. Cell viability reduction is presented in comparison to the corresponding untreated control. A linear regression model was applied for the articles reporting quality correlation analysis. The reduced number of included articles and effects for glucose level analysis was a result of non-reporting. SD = standard deviation; CI = confidence interval (with significance level of alpha = .05); LL = lower limit; UL = upper limit. *: The *p* value indicates whether there was a significant difference for the parameter phenotypes effect estimate in comparison to the effect estimate in the high glucose group.

| Moderator | Effects | Articles | Cell viability<br>reduction | SD | CI LL | CI UL | *p* value |
| --- | --- | --- | --- | --- | --- | --- | --- |
| **Glucose level** |  |  |  |  |  |  |  |
| High glucose (4500 mg/dl) | 168 | 21 | 23.1% | 3.6% | 15.8% | 30.5% | < 0.001 |
| Low glucose (1000 mg/dl) | 37 | 2 | 20.6% | 5.8% | -25.2% | 66.3% | .726* |
| No glucose | 1 | 1 | 29.0% | 3.5% | 21.6% | 36.4% | .113* |
| Not reported | 438 | 77 | 37.1% | 4.2% | 28.6% | 45.6% | .002* |
| **Articles reporting quality** |  |  |  |  |  |  |  |
| Intercept | 644 | 101 | 61.3% | 12.0% | 37.0% | 85.7% | < .001 |
| Change per unit increase<br>in the number of reported<br>parameters |  |  | -3.0% | 1.3% | -5.5% | -0.4% | .026 |

Articles reporting quality showed a significant correlation with the reported effect of TMZ (marginal *R*^*2*^ = 3.3%). Reported effect on cell viability fell by 3.0% for each unit increase in the number of reported parameters (*p* = .026). Adding these two features (glucose level and articles reporting quality) to the multivariable between-articles-variance meta-regression resulted in a slightly improved model, reducing tau^2^ from 3.8% to 3.6%.

## Discussion

### Reporting and experimental parameters

We found a highly significant relationship between the concentration of TMZ used, the duration of treatment and the measured effect of TMZ on the viability of U-87 MG cells (*p* < .001). However, the reporting of experimental parameters in this literature – including such fundamental issues as the concentration of the drug and the treatment duration – is limited. A recent study has also identified suboptimal reporting of basic experimental parameters and varying cell viability reducing effects of TMZ in *in vitro* glioma cell line experiments with TMZ single treatment (Poon et al., 2021).

Based on TMZ concentrations found in peritumoral tissue (Jackson et al., 2018; Portnow et al., 2009) and cerebrospinal fluid (Ostermann et al., 2004), Stepanenko and Checkhonin recommended *in vitro* studies should use concentrations of 1 to 10 µM (Stepanenko & Chekhonin, 2019). Although the effect of higher concentrations could be used, it seems reasonable to expect that publications use at least one clinically relevant drug concentration. More than two thirds of articles (70 of 101 articles included in meta-analysis) did not use clinically relevant TMZ concentrations in at least one of their experiments. The effects of TMZ are due to DNA alkylation and methylation, and this requires TMZ internalization, which usually occurs during cell division. An effect of TMZ can therefore be expected at the earliest after 1.5 cell doubling times, and the cell doubling time of U-87 MG is around 34 hours (ATCC, PBCF & PS-OC Bioresource Core Facility, 2012; Weller et al., 1998). Applying an early credible limit for efficacy of 51 hours, 31.7% of the articles included in the meta-analysis only measured effects before they could reasonably be expected to occur. This could lead to an underestimation of the effect of TMZ, or an overestimated effect of new drug candidates compared to TMZ, if the new drug candidates have an earlier onset of effect.

Limited reporting of key statistical properties such as the number of experimental units included (not reported in 24 of 137 articles) or the type of error presented in results (not reported in 17 articles) is of concern, and we note recommendations for improved reporting of such items (Macleod et al., 2021).

To introduce sufficient independence between repetitions of cell culture experiments, it has been suggested that experiments should be conducted over several days, with freshly prepared materials, and that the experimental unit defined as the day, so n is taken as the number of days (C. Emmerich, 2016; Lazic et al., 2018). Along with the limited reporting of the number of independent experiments and (technical) replications per experiment, this leads us to encourage researchers to clearly describe their methods for introducing robust independence.

Despite the known problems with the provenance of this cell line (Allen et al., 2016) and the widely recommended implementation of cell line authentication (Capes-Davis et al., 2019; International Cell Line Authentication Committee, 2012), we were surprised that the great majority (90.5%) of included articles did not report such an identification procedure. In addition, infrequent reporting of U-87 MG cell passage number and cell concentration used in the model adds to concern that published *in vitro* glioma research may be particularly confounded, as it is known that both parameters are potent drivers for heterogeneity (Gülden et al., 2010; OECD, 2018). Importantly, our analysis identified an encouraging but slow trend of improving reporting over time. This is in line with recent findings of a large study showing that methodological reporting quality for 1,578,964 PubMedCentral articles had increased in recent times (Menke et al., 2020). Interestingly, although showing statistical significance, the positive correlation of elevated impact factor of the journals the articles are published in and reporting quality was limited, again consistent with the findings of Menke et al. We note also that the anti-cell growth effect of TMZ was greater in publications which had less complete reporting of experimental details, consistent with previous work on *in vivo* studies which showed a higher effect of a therapeutic intervention reported in articles with worse methodological reporting and higher risks of bias (Macleod et al., 2008).

While rigid compliance to reporting guidelines may be seen by some as unduly burdensome, our findings suggest that adoption of guidelines for the design, conduct, analysis and reporting of *in vitro* research such as the MDAR framework (Macleod et al., 2021) would lead to more useful *in vitro* research.

### Sources and amplitudes of heterogeneity

The observed heterogeneity was far in excess of that expected from random sampling error (*p* < .001), even though we had taken steps to include broadly similar studies (outcome measure, culture medium). As the mean effect estimate of TMZ - across all articles with all applied drug concentrations and treatment durations – is a cell viability reduction of 33.8% compared to the untreated control, the magnitude of the SD of the effects across the articles with +/- 16.6% is almost half as high as the effect estimate itself. These strongly heterogeneous findings are in line with earlier quantifications of results repeatability in cancer research (Begley & Ellis, 2012; Prinz et al., 2011) and with the “Reproducibility Project: Cancer Biology” (eLife sciences, 2014). These investigations evaluated reproducibility in a one-on-one replication attempt and calculated the share of reproducible articles as the measure of reproducibility in a field. In contrast, our meta-analysis retrospectively extracted every published effect of a commonly performed experiment and calculated the variance of the effects across the articles as a measure of irreproducibility. We believe that this strategy has an advantage in that it enables us to recognise which study parameters may act as drivers of irreproducibility. Further, we think our approach has broader relevance, since most studies are carried out relatively independent of each other and are not the focus of one-to-one replication of selected previously published results.

The effect of TMZ was almost forty percent lower in the high glucose group than in the articles with an unreported glucose level. It is known that glucose restriction leads to a reduced cell proliferation (Bao et al., 2019) and to a sensitisation of glioblastoma cells to TMZ (Safdie et al., 2012; Wang et al., 2018), so we consider it likely that the articles with unreported glucose levels mainly used low glucose levels. However, as only two articles included in the meta-analysis reported the use of low glucose levels, we were not able to obtain precise enough estimates for the effect in the low glucose group. For the other experimental parameters tested in meta-regressions no significant moderation was observed. The results do not demonstrate that there was no effect on reproducibility and may simply reflect the statistical power of our approach depending on a minimum frequency of reporting of a particular parameter tested for its impact on reproducibility in a meta-regression. For example, only eleven studies could be included in the meta-regression analysing the relevance of different levels of cell culture confluence at time of cell passaging, and we acknowledge this is a limitation of our study.

### Implications for future research

Despite the existence of *in vitro* cell culture based experimental guidelines like the Guidance on Good Cell Culture Practice (OECD, 2018) there are no widely applied reporting guidelines specific to *in vitro* preclinical research, although the MDAR framework includes *in vitro* research. Randomised group allocation in experimental design, blinded outcome assessment and sample size calculation are well established methods to reduce risks of bias in clinical and *in vivo* research, but in this review none of the included articles reported any one of these methods. This is consistent with previous findings (C. H. Emmerich et al., 2020; The NPQIP Collaborative group, 2019), and some have argued for the implementation of randomization and blinding in *in vitro* trials (Begley & Ellis, 2012; Krithikadatta et al., 2014). Care would be required to mitigate any additional risks due to pipetting errors (because of complex pipetting schemes) and more challenging data transfers (OECD, 2018), but the risk of unconscious systematic bias in cell plating and pipetting on multi-well plates might decrease (Niepel, 2019). Although, we recommend random allocation of wells to experimental groups and the blinding of cell culture procedures and assessment of cell viability to reduce potential bias. Meaningful sample size calculations will require better understanding of the experimental unit, and we endorse the suggestion that sufficient independence between replicates should be introduced by performing experiments on different days with freshly prepared cells and reagents (Cumming et al., 2007; C. Emmerich, 2016; Lazic et al., 2018).

The choice of control in cell culture drug response assays is important, and we were concerned that the exact condition of the untreated control arm was not reported in the majority of cases. Of note, caution is advised in the selection of maximum volume percentages of common control treatments, i.e., dimethyl sulfoxide, as elevated dosage causes inhibition of cell proliferation, exposing risk for efficacy normalization of an investigated intervention (OECD, 2018). Where TMZ had been used as an active control for the evaluation of new therapeutic candidates, the parameters for the new drug were often much more detailed described than those for the control treatment with TMZ. As an example of why this might be important, if the effects of different drugs are differentially sensitive to glucose levels this may lead to erroneous interpretations of the potency of a new investigational drug.

## Limitations

Our study has several limitations. We did not choose the parameters contributing to the reporting quality based on a pre-existing reporting guideline because we could not identify an appropriate guideline for this type of experiment. Instead, we used parameters derived from previous work and from laboratory experience. It is unlikely that the chosen parameters have the same impact on heterogeneity but we had no basis to assert their different impact, and so used an unweighted score.

The overwhelming part of heterogeneity of results across the articles (89.1%) remained unexplained. As discussed earlier, the main limiting factor during the analysis was the surprisingly low frequency of reporting of experimental parameters of interest which will have limited the power of our analysis. Further reasons for unexplained heterogeneity include contributions to irreproducibility by parameters not included in the review, or different behaviours of U-87 MG cells in different laboratories for reasons unrelated to study design. Although our study included analysis on reporting of the source, authentication and cell passage, we are aware that a larger analysis with multiple different cancer cell models would be required to draw firmer conclusions. We think that the similar use of other glioblastoma cell lines means that our findings are probably transferable to these models.

Finally, we had planned to conduct a parallel review of research using more contemporary glioblastoma *in vitro* models, such as 3D glioma stem-like cells approaches (e.g. NCH421k (Campos et al., 2010)). However, in preliminary searches we were surprised to find that there was no commonly used GSC model, with authors generally using individually generated cell lines. It seemed not reasonable to perform a meta-analysis on these limited data. However, as these GSC models are probably better representatives of disease pathophysiology (Lottaz et al., 2010; Tian et al., 2017), accurate and comprehensive reporting on experimental parameters may be even more important to ensure reproducibility of their results, as elevated genetic and cellular complexity of these cells may translate into larger intrinsic biological variations.

## Conclusion

*In vitro* glioma research suffers from insufficient reporting of methods and experimental design. We believe current publication practices contribute as one source of variance that may be a driver for poor reproducibility. Although our analysis contrasts current practice with an idealised scenario and must be considered with caution in some regards (i.e., risks of bias), our study clearly supports the establishment of consensus reporting guidelines for *in vitro* (cancer) research. Our study should be considered as an independent confirmatory study of earlier reporting and reproducibility enhancing recommendations (OECD, 2018) with the additional benefits of evidence for insufficient methodological reporting as well as quantification of the reproducibility of results in a highly relevant area of *in vitro* brain cancer research. It may be relevant to raise further awareness in a wide audience of stakeholders in biomedical research.

## Acknowledgements

We thank Renfei Du (Department of Neurosurgery, University of Duesseldorf, Germany) for his assistance in literature screening and data extraction. Moreover, we are grateful to Shinichi Nakagawa (Earth and Environmental Sciences, University of New South Wales, Sydney, Australia), Igor Fischer (Department of Neurosurgery, University of Duesseldorf, Germany) and Holger Schwender (Institute of Mathematics, University of Duesseldorf, Germany) for their comments on the (meta-analytical) statistics applied.

We declare no competing interests.

## Abbreviations

ATCC: American Type Culture Collection, Manassas, Virginia
CI: Confidence interval
DMEM: Dulbecco’s Modified Eagle Medium
JIF: Journal impact factor
MTT: 3-(4,5-dimethylthiazol-2-yl)-2,5-diphenyltetrazolium bromide
SD: Standard deviation
SEM: Standard error of the mean
SyRF: Systematic Review Facility
TMZ: Temozolomide
U-87 MG: Uppsala-87 Malignant Glioma

## Notes

https://osf.io/9k3dq

https://github.com/TimoSander/Reporting-practices-as-a-source-of-heterogeneity-in-in-vitro-cancer-research/

### Competing Interest Statement

The authors have declared no competing interest.

