## Supplements 1 - 8 for "Meta-analysis on reporting practices as a source of heterogeneity in *in vitro* cancer research"

**Supplement 1:** ***Exact search strategy***

*(glioblastom* OR Astrocytom*, Grade IV OR Glioblastom* Multiform* OR Giant Cell Glioblastom* OR Brain cance* OR Malignant Gliom*) AND (Temozolomide OR Temodal OR Temodar OR methazolastone OR tmz) AND (U87 OR U-87 OR U87MG OR U87-MG OR U 87-MG OR U-87-MG OR U87 GBM OR U87-GBM OR U 87 GBM OR U-87 GBM OR U-87-GBM)*

**Supplement 2:** ***Literature screening criteria***

| Inclusion criteria | Exclusion criteria |
|---|---|
| U-87 MG cell line as glioblastoma *in vitro* model | Other models than U-87 MG cell line<br>*In vivo* models, Xenotransplantation models |
| TMZ single treatment | TMZ as a part of a combined treatment with other drugs or genetic interventions |
| Comparison of the effect of TMZ to an untreated control | No comparison to an untreated control |
| Cell viability assessment (MTT and similar colorimetric assays, cell counting) to quantify the effect of TMZ | Effect of TMZ measured with none of these cell viability assessment methods |
| DMEM as the cell culture medium | Other cell culture media than DMEM |
| Original peer-reviewed research articles | Other publication types (e.g., conference abstracts, poster presentations) |
| English language | Other languages than English |

**Suppl. 2:** Articles were included if they met all inclusion criteria and no exclusion criteria. If an article included multiple experiments where one or more experiment did not match the criteria but at least one did match, then the article was included. DMEM = Dulbecco's Modified Eagle Medium; MTT = 3-(4,5-dimethylthiazol-2-yl)-2,5-diphenyltetrazolium bromide; TMZ = temozolomide; U-87 MG = Uppsala-87 Malignant Glioma.

**Supplement 3:** ***Extracted parameters from the included articles***

| Parameter | Possible phenotypes |
|---|---|
| **General article information** | |
| Title, Authors, Year, Publishing journals JIF | n. a. |
| Conflicts of interest's statement | Conflicts / no conflicts |
| **Cell model** | |
| U-87 MG cell line source | Name and country of source |
| U-87 MG cell line authentication reporting | Reported / not reported |
| U-87 MG cell line age reporting | Reported / not reported |
| **Cell culture conditions** | |
| Number/Volume/Concentration of U-87 MG cells | Exact Number/Volume/Concentration |
| Cell passaging criterion | Confluency/Time intervals |
| Glucose level of cell culture medium | Exact glucose concentration / high glucose / low glucose / no glucose |
| Reporting of successful mycoplasma exclusion test | Reported / not reported |
| Supplemented antibiotics | Name and dose of the antibiotics |
| FBS use and source | FBS supplemented / FBS not supplemented |
| | Name and country of source |
| **Intervention & control group** | |
| TMZ source | Name and country of source |
| Volume of the added TMZ suspension | Exact volume |
| Type of untreated control | Drug vehicle / medium only / other |
| Volume of the added control suspension | Exact volume |
| **Outcome** | |
| Cell viability of U-87 MG cells after incubation with TMZ and the corresponding untreated control | Mean and error data of cell viabilities / growth inhibition rates / proliferation rates |
| Concentration of TMZ | Exact concentration |
| Treatment duration | Exact duration |
| Number of experiments | Exact number of experiments |
| Type of outcome measurement assay | Name of assay |

**Suppl. 3:** FBS = Fetal bovine serum; JIF = Journal Impact Factor (for the year of publication; obtained from Clarivates InCites Journal Citation Reports); U-87 MG = Uppsala-87 Malignant Glioma; TMZ = Temozolomide.

**Supplement 4:** ***Assessed potential risks of bias parameters***

| |
|---|
| Were data for every relevant experiment mentioned in the methods section of a particular article presented? |
| Were data for every concentration of temozolomide and treatment duration presented? |
| Were the number of experiments and the number of replicates per experiment clearly reported? |
| Was a sample size calculation for the needed number of experiments reported? |
| Was the way calculating the U87-MG cell viability mean and error data clearly reported? |
| Was it reported whether the authors allocated the U87-MG cells randomly to the treatment and control group? |
| Was it reported whether the authors measured the U87-MG cell viability blinded? |
| Was a pre-registered study protocol available? |

**Suppl. 4:** U-87 MG = Uppsala-87 Malignant Glioma.

**Supplement 5:** ***Parameters the authors were asked for if they were not reported in the article (and additional potential risks of bias parameters)***

| |
|---|
| *Missing information in the articles:* |
| U-87 MG cell line authentication |
| U-87 MG age (maximum number of passages) |
| Glucose level of cell culture medium |
| Type of untreated control |
| Concentration of TMZ |
| Number of experiments and replicates per experiment |
| Type of error of the presented data (SD or SEM) |
| Treatment duration |
| *Additional risks of bias parameters:* |
| Therapy regime (single dose, multi dose or different) |
| Was the cell viability measured directly after the treatment of U-87 MG cells with TMZ? |
| Way of calculation of cell viability data (mean of all data, mean of means or different) |

**Suppl. 5:** SD = standard deviation; SEM = standard error of the mean; U-87 MG = Uppsala-87 Malignant Glioma; TMZ = Temozolomide.

**Supplement 6: Full list of *included articles into the systematic review and meta-analysis***

| Article number | Title | Authors | Year | DOI | Included in meta-analysis |
|---|---|---|---|---|---|
| 1 | Afatinib and Temozolomide combination inhibits tumorigenesis by targeting EGFRvIII-cMet signaling in glioblastoma cells | Vengoji et al. | 2019 | 10.1186/s13046-019-1264-2 | TRUE |
| 2 | Akt and beta-catenin contribute to TMZ resistance and EMT of MGMT negative malignant glioma cell line | Yi et al. | 2016 | 10.1016/j.jns.2016.05.054 | TRUE |
| 3 | Anti-tumor activities of luteolin and silibinin in glioblastoma cells: overexpression of miR-7-1-3p augmented luteolin and silibinin to inhibit autophagy and induce apoptosis in glioblastoma in vivo | Chakrabarti et al. | 2015 | 10.1007/s10495-015-1198-x | TRUE |
| 4 | Anticancer activity of flavonoids isolated from Achyrocline satureioides in gliomas cell lines | De Souza et al. | 2018 | 10.1016/j.tiv.2018.04.013 | TRUE |
| 5 | Artesunate Enhances the Antiproliferative Effect of Temozolomide on U87MG and A172 Glioblastoma Cell Lines | Karpel-Massler et al. | 2014 | 10.2174/18715206113136660340 | TRUE |
| 6 | ATM inhibitor KU-55933 increases the TMZ responsiveness of only inherently TMZ sensitive GBM cells | Nadkarni et al. | 2012 | 10.1007/s11060-012-0979-0 | TRUE |
| 7 | Berberine induces senescence of human glioblastoma cells by downregulating the EGFR-MEK-ERK signaling pathway | Liu et al. | 2014 | 10.1158/1535-7163.MCT-14-0634 | TRUE |
| 8 | beta-elemene enhances both radiosensitivity and chemosensitivity of glioblastoma cells through the inhibition of the ATM signaling pathway | Liu et al. | 2015 | 10.3892/or.2015.4050 | TRUE |
| 9 | Blocking LDHA glycolytic pathway sensitizes glioblastoma cells to radiation and temozolomide | Koukourakis et al. | 2017 | 10.1016/j.bbrc.2017.07.138 | TRUE |
| 10 | Bortezomib inhibits growth and sensitizes glioma to temozolomide (TMZ) via down-regulating the FOXM1-Survivin axis | Tang et al. | 2019 | 10.1186/s40880-019-0424-2 | TRUE |

| | | | | | |
|---|---|---|---|---|---|
| 11 | Bortezomib overcomes MGMT-related resistance of glioblastoma cell lines to temozolomide in a schedule-dependent manner | Vlachostergios et al. | 2013 | 10.1007/s10637-013-9968-1 | TRUE |
| 12 | Bufothionine Promotes Apoptosis via Triggering ER Stress and Synergizes with Temozolomide in Glioblastoma Multiforme Cells | Sun et al. | 2019 | 10.1002/ar.24194 | TRUE |
| 13 | Calpain suppresses cell growth and invasion of glioblastoma multiforme by producing the cleavage of filamin A | Cai et al. | 2020 | 10.1007/s10147-020-01636-7 | TRUE |
| 14 | Chemotherapeutic effect of tamoxifen on temozolomide-resistant gliomas | He et al. | 2015 | 10.1097/CAD.0000000000000197 | TRUE |
| 15 | Chloroquine enhances temozolomide cytotoxicity in malignant gliomas by blocking autophagy | Golden et al. | 2014 | 10.3171/2014.9.FOCUS14504 | TRUE |
| 16 | Chronic exposure of human glioblastoma tumors to low concentrations of a pesticide mixture induced multidrug resistance against chemotherapy agents | Doganlar et al. | 2020 | 10.1016/j.ecoenv.2020.110940 | TRUE |
| 17 | Combination of biochanin a and temozolomide impairs tumor growth by modulating cell metabolism in glioblastoma multiforme | Desai et al. | 2019 | 10.21873/anticanres.13079 | TRUE |
| 18 | Combination of caspase transfer using the human telomerase reverse transcriptase promoter and conventional therapies for malignant glioma cells | Takeuchi et al. | 2004 | 10.3892/ijo.25.1.57 | TRUE |
| 19 | Combination of the mTOR inhibitor RAD001 with temozolomide and radiation effectively inhibits the growth of glioblastoma cells in culture | Burckel et al. | 2014 | 10.3892/or.2014.3590 | TRUE |
| 20 | Combined effects of mesenchymal stem cells carrying cytosine deaminase gene with 5-fluorocytosine and temozolomide in orthotopic glioma model | Chang et al. | 2020 | | TRUE |

| | | | | | |
|---|---|---|---|---|---|
| 21 | CtIP contributes to non-homologous end joining formation through interacting with ligase IV and promotion of TMZ resistance in glioma cells | Yang et al. | 2019 | 10.26355/eurrev_201903_17252 | TRUE |
| 22 | Cytotoxicity of temozolomide on human glioblastoma cells is enhanced by the concomitant exposure to an extremely low-frequency electromagnetic field (100 Hz 100 G) | Akbarnejad et al. | 2017 | 10.1016/j.biopha.2017.05.050 | TRUE |
| 23 | Development of transferrin-modified poly(lactic-co-glycolic acid) nanoparticles for glioma therapy | Mao et al. | 2019 | 10.1097/CAD.0000000000000754 | TRUE |
| 24 | Do Anti-Oxidants Vitamin D(3) Melatonin and Alpha-Lipoic Acid Have Synergistic Effects with Temozolomide on Cultured Glioblastoma Cells? | McConnell et al. | 2018 | 10.3390/medicines5020058 | TRUE |
| 25 | Down-Regulation of AQP4 Expression via p38 MAPK Signaling in Temozolomide-Induced Glioma Cells Growth Inhibition and Invasion Impairment | Chen et al. | 2017 | 10.1002/jcb.26176 | TRUE |
| 26 | Effect of the STAT3 inhibitor STX-0119 on the proliferation of a temozolomide-resistant glioblastoma cell line | Ashizawa et al. | 2014 | 10.3892/ijo.2014.2439 | TRUE |
| 27 | Effects of solvent used for fabrication on drug loading and release kinetics of electrosprayed temozolomide-loaded PLGA microparticles for the treatment of glioblastoma | Rodriguez de Anda et al. | 2019 | 10.1002/jbm.b.34324 | TRUE |
| 28 | Effects of temozolomide (TMZ) on the expression and interaction of heat shock proteins (HSPs) and DNA repair proteins in human malignant glioma cells | Castro et al. | 2014 | 10.1007/s12192-014-0537-0 | TRUE |
| 29 | EGCG inhibits properties of glioma stem-like cells and synergizes with temozolomide through downregulation of P-glycoprotein inhibition | Zhang et al. | 2014 | 10.1007/s11060-014-1604-1 | TRUE |

| | | | | | |
|---|---|---|---|---|---|
| 30 | EMAP-II sensitize U87MG and glioma stem-like cells to temozolomide via induction of autophagy-mediated cell death and G2/M arrest | Yu et al. | 2017 | 10.1080/15384101.2017.1315492 | TRUE |
| 31 | Enhancing glioblastoma cell sensitivity to chemotherapeutics: A strategy involving survivin gene silencing mediated by gemini surfactant-based complexes | Cruz et al. | 2016 | 10.1016/j.ejpb.2016.04.014 | TRUE |
| 32 | Exogenous IGFBP-2 promotes proliferation invasion and chemoresistance to temozolomide in glioma cells via the integrin beta 1-ERK pathway | Han et al. | 2014 | 10.1038/bjc.2014.435 | TRUE |
| 33 | Fever-Range Hyperthermia vs. Hypothermia Effect on Cancer Cell Viability Proliferation and HSP90 Expression | Kalamida et al. | 2015 | 10.1371/journal.pone.0116021 | TRUE |
| 34 | FTY720 inhibits the Nrf2/ARE pathway in human glioblastoma cell lines and sensitizes glioblastoma cells to temozolomide | Zhang et al. | 2017 | 10.1016/j.pharep.2017.07.003 | TRUE |
| 35 | G3BP1 knockdown sensitizes U87 glioblastoma cell line to Bortezomib by inhibiting stress granules assembly and potentializing apoptosis | Bittencourt et al. | 2019 | 10.1007/s11060-019-03252-6 | TRUE |
| 36 | GADD45A plays a protective role against temozolomide treatment in glioblastoma cells | Wang et al. | 2017 | 10.1038/s41598-017-06851-3 | TRUE |
| 37 | Gene expression profiling predicts response to temozolomide in malignant gliomas | Yoshino et al. | 2010 | 10.3892/ijo_00000621 | TRUE |
| 38 | Genomic profiling of long non-coding RNA and mRNA expression associated with acquired temozolomide resistance in glioblastoma cells | Zheng et al. | 2017 | 10.3892/ijo.2017.4033 | TRUE |
| 39 | Glucosylceramide synthase silencing combined with the receptor tyrosine kinase inhibitor axitinib as a new multimodal strategy for glioblastoma | Morais et al. | 2019 | 10.1093/hmg/ddz152 | TRUE |

| | | | | | |
|---|---|---|---|---|---|
| 40 | Growth Inhibitory Effects of Dipotassium Glycyrrhizinate in Glioblastoma Cell Lines by Targeting MicroRNAs Through the NF-kappa B Signaling Pathway | Bonafe et al. | 2019 | 10.3892/ijmm.2015.2312 | TRUE |
| 41 | Heterogeneous glioblastoma cell cross-talk promotes phenotype alterations and enhanced drug resistance | Motaln et al. | 2015 | 10.18632/oncotarget.5701 | TRUE |
| 42 | High-throughput screening uncovers miRNAs enhancing glioblastoma cell susceptibility to tyrosine kinase inhibitors | Cunha et al. | 2017 | 10.1093/hmg/ddx323 | TRUE |
| 43 | Honokiol enhances temozolomide-induced apoptotic insults to malignant glioma cells via an intrinsic mitochondrion-dependent pathway | Chio et al. | 2018 | 10.1016/j.phymed.2018.06.012 | TRUE |
| 44 | IDH1 R132H mutation regulates glioma chemosensitivity through Nrf2 pathway | Li et al. | 2017 | 10.18632/oncotarget.15868 | TRUE |
| 45 | Improved effects of honokiol on temozolomide-induced autophagy and apoptosis of drug-sensitive and -tolerant glioma cells | Chio et al. | 2018 | 10.1186/s12885-018-4267-z | TRUE |
| 46 | In vitro and in vivo effect of human lactoferrin on glioblastoma growth | Arcella et al. | 2015 | 10.3171/2014.12.JNS14512 | TRUE |
| 47 | In vitro novel combinations of psychotropics and anti-cancer modalities in U87 human glioblastoma cells | Tzadok et al. | 2010 | 10.3892/ijo-00000756 | TRUE |
| 48 | In vitro radiosensitizing effects of temozolomide on U87MG cell lines of human glioblastoma multiforme | Borhani et al. | 2017 | | TRUE |
| 49 | Induction of microRNA-146a is involved in curcumin-mediated enhancement of temozolomide cytotoxicity against human glioblastoma | Wu et al. | 2015 | 10.3892/mmr.2015.4087 | TRUE |
| 50 | Inhibition of STAT3 reverses alkylator resistance through modulation of the AKT and beta-catenin signaling pathways | Wang et al. | 2011 | 10.3892/or.2011.1396 | TRUE |

| | | | | | |
|---|---|---|---|---|---|
| 51 | Inhibition of telomerase activity in malignant glioma cells correlates with their sensitivity to temozolomide | Kanzawa et al. | 2003 | 10.1038/sj.bjc.6601193 | TRUE |
| 52 | Lithium enhances the antitumour effect of temozolomide against TP53 wild-type glioblastoma cells via NFAT1/FasL signalling | Han et al. | 2017 | 10.1038/bjc.2017.89 | TRUE |
| 53 | Magnolol and honokiol exert a synergistic anti-tumor effect through autophagy and apoptosis in human glioblastomas | Cheng et al. | 2016 | 10.18632/oncotarget.8674 | TRUE |
| 54 | Major Contribution of Caspase-9 to Honokiol-Induced Apoptotic Insults to Human Drug-Resistant Glioblastoma Cells | Wu et al. | 2020 | 10.3390/molecules25061450 | TRUE |
| 55 | Mechanisms and antitumor activity of a binary EGFR/DNA-targeting strategy overcomes resistance of glioblastoma stem cells to temozolomide | Sharifi et al. | 2019 | 10.1158/1078-0432.CCR-19-0955 | TRUE |
| 56 | MicroRNA-182 targets protein phosphatase 1 regulatory inhibitor subunit 1C in glioblastoma | Liu et al. | 2017 | 10.18632/oncotarget.21309 | TRUE |
| 57 | MicroRNA-21 silencing enhances the cytotoxic effect of the antiangiogenic drug sunitinib in glioblastoma | Costa et al. | 2013 | 10.1093/hmg/dds496 | TRUE |
| 58 | MicroRNA-29b promotes cell sensitivity to Temozolomide by targeting STAT3 in glioma | Xu et al. | 2020 | 10.26355/eurrev_202002_20370 | TRUE |
| 59 | MIM1 the Mcl-1 - specific BH3 mimetic induces apoptosis in human U87MG glioblastoma cells | Respondek et al. | 2018 | 10.1016/j.tiv.2018.08.007 | TRUE |
| 60 | MiR-144 overexpression as a promising therapeutic strategy to overcome glioblastoma cell invasiveness and resistance to chemotherapy | Cardoso et al. | 2019 | 10.1093/hmg/ddz099 | TRUE |
| 61 | miR-203 sensitizes glioma cells to temozolomide and inhibits glioma cell invasion by targeting E2F3 | Tang et al. | 2015 | 10.3892/mmr.2014.3101 | TRUE |
| 62 | MiR-519a enhances chemosensitivity and promotes autophagy in glioblastoma by targeting STAT3/Bcl2 signaling | Li et al. | 2018 | 10.1186/s13045-018-0618-0 | TRUE |

| | | | | | |
|---|---|---|---|---|---|
| | pathway | | | | |
| 63 | Mitochondria Transcription Factor A: A Putative Target for the Effect of Melatonin on U87MG Malignant Glioma Cell Line | Franco et al. | 2018 | 10.3390/molecules 23051129 | TRUE |
| 64 | N-(2-hydroxyphenyl)acetamide (NA-2) and Temozolomide synergistically induce apoptosis in human glioblastoma cell line U87 | Hanif et al. | 2014 | 10.1186/s12935-014-0133-5 | TRUE |
| 65 | Polyphyllin VII Promotes Apoptosis and Autophagic Cell Death via ROS-Inhibited AKT Activity and Sensitizes Glioma Cells to Temozolomide | Pang et al. | 2019 | 10.1155/2019/1805 635 | TRUE |
| 66 | Quercetin sensitizes human glioblastoma cells to temozolomide in vitro via inhibition of Hsp27 | Sang et al. | 2014 | 10.1038/aps.2014.2 2 | TRUE |
| 67 | Radiobiological evaluation and correlation with the local effect model (LEM) of carbon ion radiation therapy and temozolomide in glioblastoma cell lines | Combs et al. | 2008 | 10.1080/09553000 802641151 | TRUE |
| 68 | Receptor-mediated PLGA nanoparticles for glioblastoma multiforme treatment | Ramalho et al. | 2018 | 10.1016/j.ijpharm.2 018.04.062 | TRUE |
| 69 | Regulation of Integrated Stress Response Sensitizes U87MG Glioblastoma Cells to Temozolomide Through the Mitochondrial Apoptosis Pathway | He et al. | 2018 | 10.1002/ar.23839 | TRUE |
| 70 | Riluzole enhances the antitumor effects of temozolomide via suppression of MGMT expression in glioblastoma | Yamada et al. | 2020 | 10.3171/2019.12.J NS192682 | TRUE |
| 71 | Salvianolic acid B renders glioma cells more sensitive to radiation via Fis-1-mediated mitochondrial dysfunction | Chen et al. | 2018 | 10.1016/j.biopha.20 18.08.113 | TRUE |
| 72 | Sequence-dependent synergistic inhibition of human glioma cell lines by combined temozolomide and miR-21 inhibitor gene therapy | Qian et al. | 2012 | 10.1021/mp300203 9 | TRUE |

| | | | | | |
|---|---|---|---|---|---|
| 73 | Sequential treatment of phenethyl isothiocyanate increases sensitivity of temozolomide resistant glioblastoma cells by decreasing expression of mgmt via nf-kappab pathway | Guo et al. | 2019 | | TRUE |
| 74 | Silencing SATB1 overcomes temozolomide resistance by downregulating MGMT expression and upregulating SLC22A18 expression in human glioblastoma cells | Yang et al. | 2018 | 10.1038/s41417-018-0040-3 | TRUE |
| 75 | Sirtuin 1 knockdown inhibits glioma cell proliferation and potentiates temozolomide toxicity via facilitation of reactive oxygen species generation | Chen et al. | 2019 | 10.3892/ol.2019.10235 | TRUE |
| 76 | STAT3 Inhibition Overcomes Temozolomide Resistance in Glioblastoma by Downregulating MGMT Expression | Kohsaka et al. | 2012 | 10.1158/1535-7163.MCT-11-0801 | TRUE |
| 77 | Study on therapeutic action and mechanism of TMZ Combined with RITA against glioblastoma | Wu et al. | 2018 | 10.1159/000495923 | TRUE |
| 78 | Suppression of the Eag1 potassium channel sensitizes glioblastoma cells to injury caused by temozolomide | Sales et al. | 2016 | 10.3892/ol.2016.4992 | TRUE |
| 79 | Synergistic inhibition of human glioma cell line by temozolomide and PAMAM-mediated miR-21i | Qian et al. | 2012 | 10.1002/app.37823 | TRUE |
| 80 | Synergistic suppression of noscapine and conventional chemotherapeutics on human glioblastoma cell growth | QI et al. | 2013 | 10.1038/aps.2013.40 | TRUE |
| 81 | Targeted Brain Tumor Therapy by Inhibiting the MDM2 Oncogene: In Vitro and In Vivo Antitumor Activity and Mechanism of Action | Punganuru et al. | 2020 | 10.3390/cells9071592 | TRUE |
| 82 | TAZ promotes temozolomide resistance by upregulating MCL-1 in human glioma cells | Tian et al. | 2015 | 10.1016/j.bbrc.2015.05.115 | TRUE |
| 83 | Temozolomide Cocrystals Exhibit Drug Sensitivity in Glioblastoma Cells | Kusuma et al. | 2014 | 10.1007/s40010-014-0142-8 | TRUE |

| | | | | | |
|---|---|---|---|---|---|
| 84 | Temozolomide induces autophagy via ATM-AMPK-ULK1 pathways in glioma | Zou et al. | 2014 | 10.3892/mmr.2014.2151 | TRUE |
| 85 | The DNA repair protein ALKBH2 mediates temozolomide resistance in human glioblastoma cells | Johannessen et al. | 2012 | 10.1093/neuonc/nos301 | TRUE |
| 86 | The Effect of Ascorbic Acid over the Etoposide- and Temozolomide-Mediated Cytotoxicity in Glioblastoma Cell Culture: A Molecular Study | Gokturk et al. | 2018 | 10.5137/1019-5149.JTN.19111-16.1 | TRUE |
| 87 | The effect of polysaccharides from Cibotium barometz on enhancing temozolomide-induced glutathione exhausted in human glioblastoma U87 cells as revealed by H-1 NMR metabolomics analysis | Shi et al. | 2020 | 10.1016/j.ijbiomac.2020.03.243 | TRUE |
| 88 | The effect of silibinin in enhancing toxicity of temozolomide and etoposide in p53 and PTEN-mutated resistant glioma cell lines | Elhag et al. | 2015 | | TRUE |
| 89 | The Effect of Temozolomide/Poly(lactide-co-glycolide) (PLGA)/Nano-Hydroxyapatite Microspheres on Glioma U87 Cells Behavior | Zhang et al. | 2012 | 10.3390/ijms13011109 | TRUE |
| 90 | The HIV-derived protein Vpr52-96 has anti-glioma activity in vitro and in vivo | Kübler et al. | 2016 | 10.1158/1538-7445.AM2015-4458 | TRUE |
| 91 | The mTOR inhibitor RAD001 potentiates autophagic cell death induced by temozolomide in a glioblastoma cell line | Josset et al. | 2013 | | TRUE |
| 92 | The Pan-Bcl-2 Inhibitor (-)-Gossypol Triggers Autophagic Cell Death in Malignant Glioma | Voss et al. | 2010 | 10.1158/1541-7786.MCR-09-0562 | TRUE |
| 93 | The Synergistic Effect of Combination Progesterone and Temozolomide on Human Glioblastoma Cells | Atif et al. | 2015 | 10.1371/journal.pone.0131441 | TRUE |
| 94 | The synergistic effect of combination temozolomide and chloroquine treatment is dependent on autophagy formation and p53 status in glioma cells | Lee et al. | 2015 | 10.1016/j.canlet.2015.02.012 | TRUE |

| | | | | | |
|---|---|---|---|---|---|
| 95 | Tim-3 expression in glioma cells is associated with drug resistance | Zhang et al. | 2019 | 10.4103/jcrt.JCRT_630_18 | TRUE |
| 96 | Tramadol attenuates the sensitivity of glioblastoma to temozolomide through the suppression of Cx43-mediated gap junction intercellular communication | Wang et al. | 2018 | 10.3892/ijo.2017.4188 | TRUE |
| 97 | Transcriptional targeting of adenovirally delivered tumor necrosis factor alpha by temozolomide in experimental glioblastoma | Yamini et al. | 2004 | 10.1158/0008-5472.CAN-04-2117 | TRUE |
| 98 | Verapamil potentiates anti-glioblastoma efficacy of temozolomide by modulating apoptotic signaling | Hanif et al. | 2018 | 10.1016/j.tiv.2018.07.001 | TRUE |
| 99 | Verbascoside inhibits glioblastoma cell proliferation migration and invasion while promoting apoptosis through upregulation of protein tyrosine phosphatase SHP-1 and inhibition of STAT3 phosphorylation | Jia et al. | 2018 | 10.1159/000491067 | TRUE |
| 100 | Zinc enhances TMZ cytotoxicity in glioblastoma multiforme model system | Toren et al. | 2016 | 10.1093/neuonc/not176 | TRUE |
| 101 | β-Elemene inhibits proliferation through crosstalk between glia maturation factor β and extracellular signal-regulated kinase 1/2 and impairs drug resistance to temozolomide in glioblastoma cells | Zhu et al. | 2014 | 10.3892/mmr.2014.2273 | TRUE |
| 102 | Anti-epidermal growth factor receptor siRNA carried by chitosan-transacylated lipid nanocapsules increases sensitivity of glioblastoma cells to temozolomide | Messaoudi et al. | 2014 | 10.2147/IJN.S59134 | FALSE |
| 103 | Autophagic flux response and glioblastoma sensitivity to radiation | Mitrakas et al. | 2018 | 10.20892/j.issn.2095-3941.2017.0173 | FALSE |
| 104 | Autophagy mediates glucose starvation-induced glioblastoma cell quiescence and chemoresistance through coordinating cell metabolism cell | Wang et al. | 2018 | 10.1038/s41419-017-0242-x | FALSE |

| | | | | | |
|---|---|---|---|---|---|
| | cycle and survival | | | | |
| 105 | BET inhibitor I-BET151 sensitizes GBM cells to temozolomide via PUMA induction | Yao et al. | 2019 | 10.1038/s41417-018-0068-4 | FALSE |
| 106 | Combined effect of 2-5A-linked antisense against telomerase RNA and conventional therapies on human malignant glioma cells in vitro and in vivo | Iwado et al. | 2007 | 10.3892/ijo.31.5.1087 | FALSE |
| 107 | Decreasing GSH and increasing ROS in chemosensitivity gliomas with IDH1 mutation | Shi et al. | 2014 | 10.1007/s13277-014-2644-z | FALSE |
| 108 | Downregulation of Id2 increases chemosensitivity of glioma | Zhao et al. | 2015 | 10.1007/s13277-015-3055-5 | FALSE |
| 109 | Downregulation of miR-155 inhibits proliferation and enhances chemosensitivity to Temozolomide in glioma cells | Meng et al. | 2017 | | FALSE |
| 110 | Downregulation of miR-196b Promotes Glioma Cell Sensitivity to Temozolomide Chemotherapy and Radiotherapy | Ma et al. | 2018 | | FALSE |
| 111 | Effect of temozolomide on the viability of musculoskeletal sarcoma cells | Kusabe et al. | 2015 | 10.3892/ol.2015.3506 | FALSE |
| 112 | Effects of galbanic acid on proliferation migration and apoptosis of glioblastoma cells through PI3K/Akt/mTOR signaling pathway | Shahcheraghi et al. | 2020 | 10.2174/1874467213666200512075507 | FALSE |
| 113 | Encapsulation of Temozolomide in a Calixarene Nanocapsule Improves Its Stability and Enhances Its Therapeutic Efficacy against Glioblastoma | Renziehausen et al. | 2019 | 10.1158/1535-7163.MCT-18-1250 | FALSE |
| 114 | FTO inhibition enhances the anti-tumor effect of temozolomide by targeting MYC-miR-155/23a cluster-MXI1 feedback circuit in glioma | Xiao et al. | 2020 | 10.1158/0008-5472.CAN-20-0132 | FALSE |
| 115 | Green tea epigallocatechin gallate enhances therapeutic efficacy of temozolomide in orthotopic mouse | Chen et al. | 2011 | 10.1016/j.canlet.2010.11.008 | FALSE |

| | | | | | |
|---|---|---|---|---|---|
| | glioblastoma models | | | | |
| 116 | Growth-inhibitory and chemosensitizing effects of microRNA-31 in human glioblastoma multiforme cells | Zhou et al. | 2015 | 10.3892/ijmm.2015.2312 | FALSE |
| 117 | Growth-inhibitory and chemosensitizing effects of the glutathione-S- transferase-pi-activated nitric oxide donor PABA/NO in malignant gliomas | Kogias et al. | 2012 | 10.1002/ijc.26106 | FALSE |
| 118 | Identification of Key Candidate Proteins and Pathways Associated with Temozolomide Resistance in Glioblastoma Based on Subcellular Proteomics and Bioinformatical Analysis | Yi et al. | 2018 | 10.1155/2018/5238760 | FALSE |
| 119 | Inhibition of EZH2 reverses chemotherapeutic drug TMZ chemosensitivity in glioblastoma | Fan et al. | 2014 | | FALSE |
| 120 | Inhibition of JNK Potentiates Temozolomide-induced Cytotoxicity in U87MG Glioblastoma Cells via Suppression of Akt Phosphorylation | Vo et al. | 2014 | | FALSE |
| 121 | Long noncoding RNA RP11-838N2.4 enhances the cytotoxic effects of temozolomide by inhibiting the functions of miR-10a in glioblastoma cell lines | Liu et al. | 2016 | 10.18632/oncotarget.9699 | FALSE |
| 122 | LY294002 enhances cytotoxicity of temozolomide in glioma by down-regulation of the PI3K/Akt pathway | Chen et al. | 2012 | 10.3892/mmr.2011.674 | FALSE |
| 123 | Mechanisms operative in the antitumor activity of temozolomide in glioblastoma multiforme | Fischer et al. | 2007 | 10.1097/PPO.0b013e318157053f | FALSE |
| 124 | MiR-21 protected human glioblastoma U87MG cells from chemotherapeutic drug temozolomide induced apoptosis by decreasing Bax/Bcl-2 ratio and caspase-3 activity | Shi et al. | 2010 | 10.1016/j.brainres.2010.07.009 | FALSE |
| 125 | Mutant TP53 enhances the resistance of glioblastoma cells to temozolomide by up-regulating O- | Wang et al. | 2012 | 10.1007/s10072-012-1257-9 | FALSE |

| | | | | | |
|---|---|---|---|---|---|
| | 6-methylguanine DNA-methyltransferase | | | | |
| 126 | Next Generation Sequencing-Based Transcriptome Predicts Bevacizumab Efficacy in Combination with Temozolomide in Glioblastoma | Adilijiang et al. | 2019 | 10.3390/molecules24173046 | FALSE |
| 127 | Olanzapine inhibits proliferation migration and anchorage-independent growth in human glioblastoma cell lines and enhances temozolomide's antiproliferative effect | Karpel-Massler et al. | 2014 | 10.1007/s11060-014-1688-7 | FALSE |
| 128 | Ovatodiolide inhibits the oncogenicity and cancer stem cell-like phenotype of glioblastoma cells as well as potentiate the anticancer effect of temozolomide | Su et al. | 2019 | 10.1016/j.phymed.2019.152840 | FALSE |
| 129 | Overexpression of iASPP-SV in glioma is associated with poor prognosis by promoting cell viability and antagonizing apoptosis | Liu et al. | 2015 | 10.1007/s13277-015-4503-y | FALSE |
| 130 | Polymer - Temozolomide Conjugates as Therapeutics for Treating Glioblastoma | Ward et al. | 2018 | 10.1021/acs.molpharmaceut.8b00766 | FALSE |
| 131 | Potentiation of antiglioma effect with combined temozolomide and interferon-beta | Park et al. | 2006 | | FALSE |
| 132 | Silence of bFGF enhances chemosensitivity of glioma cells to temozolomide through the MAPK signal pathway | Wang et al. | 2016 | 10.1093/abbs/gmw035 | FALSE |
| 133 | Synergistic combination of chemo-phototherapy based on temozolomide/ICG-loaded iron oxide nanoparticles for brain cancer treatment | Kwon et al. | 2019 | 10.3892/or.2019.7289 | FALSE |
| 134 | Temozolomide Gemcitabine and Decitabine Hybrid Nanoconjugates: From Design to Proof-of-Concept (PoC) of Synergies toward the Understanding of Drug Impact on Human Glioblastoma Cells | Sahli et al. | 2020 | 10.1021/acs.jmedchem.0c00694 | FALSE |

| | | | | | |
|---|---|---|---|---|---|
| 135 | The synergic antitumor effects of paclitaxel and temozolomide co-loaded in mPEG-PLGA nanoparticles on glioblastoma cells | Xu et al. | 2016 | 10.18632/oncotarget.7896 | FALSE |
| 136 | Vincristine and temozolomide combined chemotherapy for the treatment of glioma: a comparison of solid lipid nanoparticles and nanostructured lipid carriers for dual drugs delivery | Wu et al. | 2015 | 10.3109/10717544.2015.1058434 | FALSE |
| 137 | YKL-40 downregulation is a key factor to overcome temozolomide resistance in a glioblastoma cell line | Akiyama et al. | 2014 | 10.3892/or.2014.3195 | FALSE |

**Suppl. 6:** Extracted data for every included article are provided in the supplementary datasets that are available on GitHub (https://github.com/TimoSander/Reporting-practices-as-a-source-of-heterogeneity-in-in-vitro-cancer-research/).

**Supplement 7:** ***Reported temozolomide concentrations in the included studies into meta-analysis***

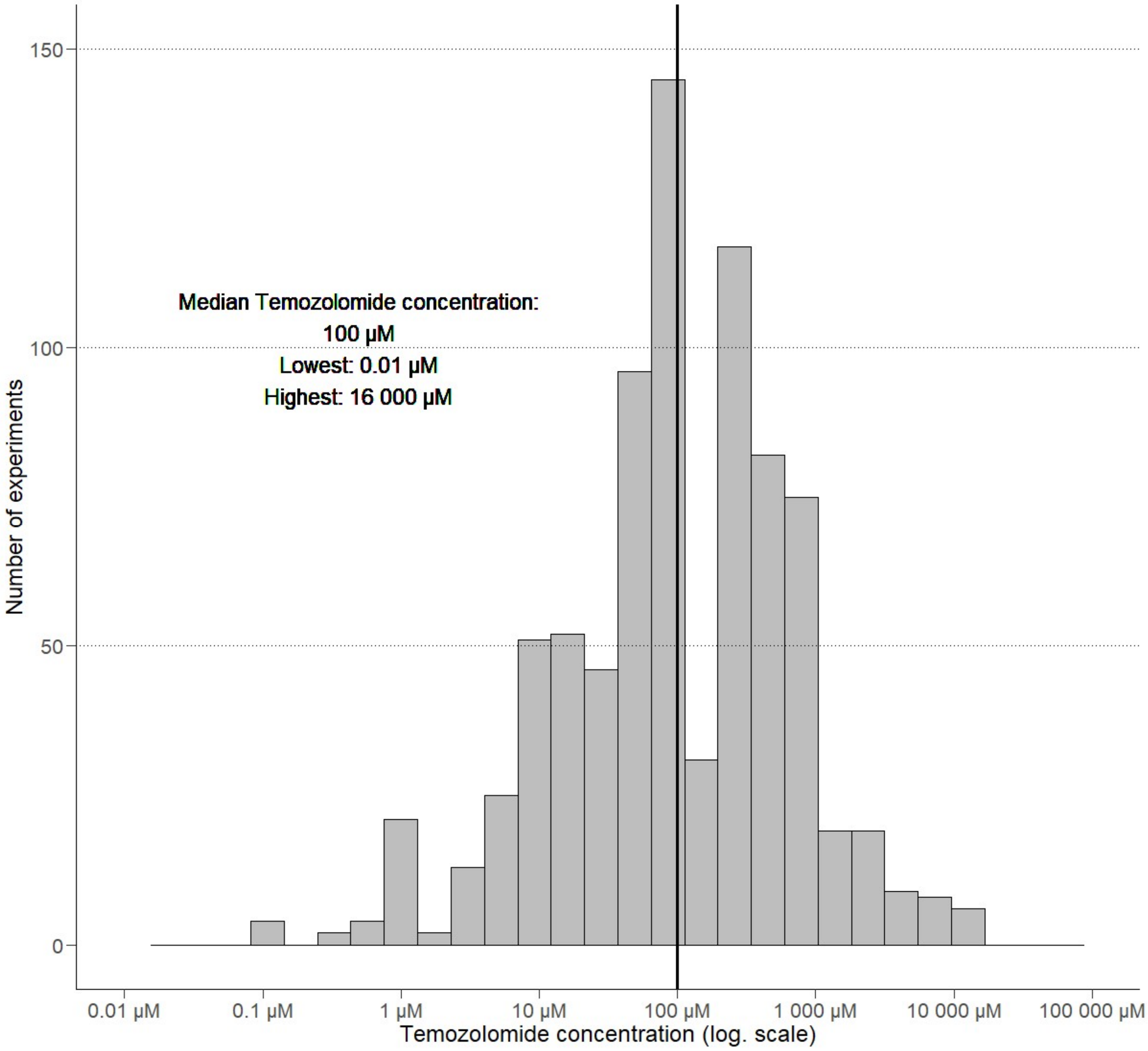


**Suppl. 7:** 137 articles included cell viability data for 828 temozolomide concentrations (98 unique). experiments using 19 treatment durations. The drug's concentration is presented on a logarithmic scale.

**Supplement 8:** ***Reported treatment durations in the included studies into meta-analysis***

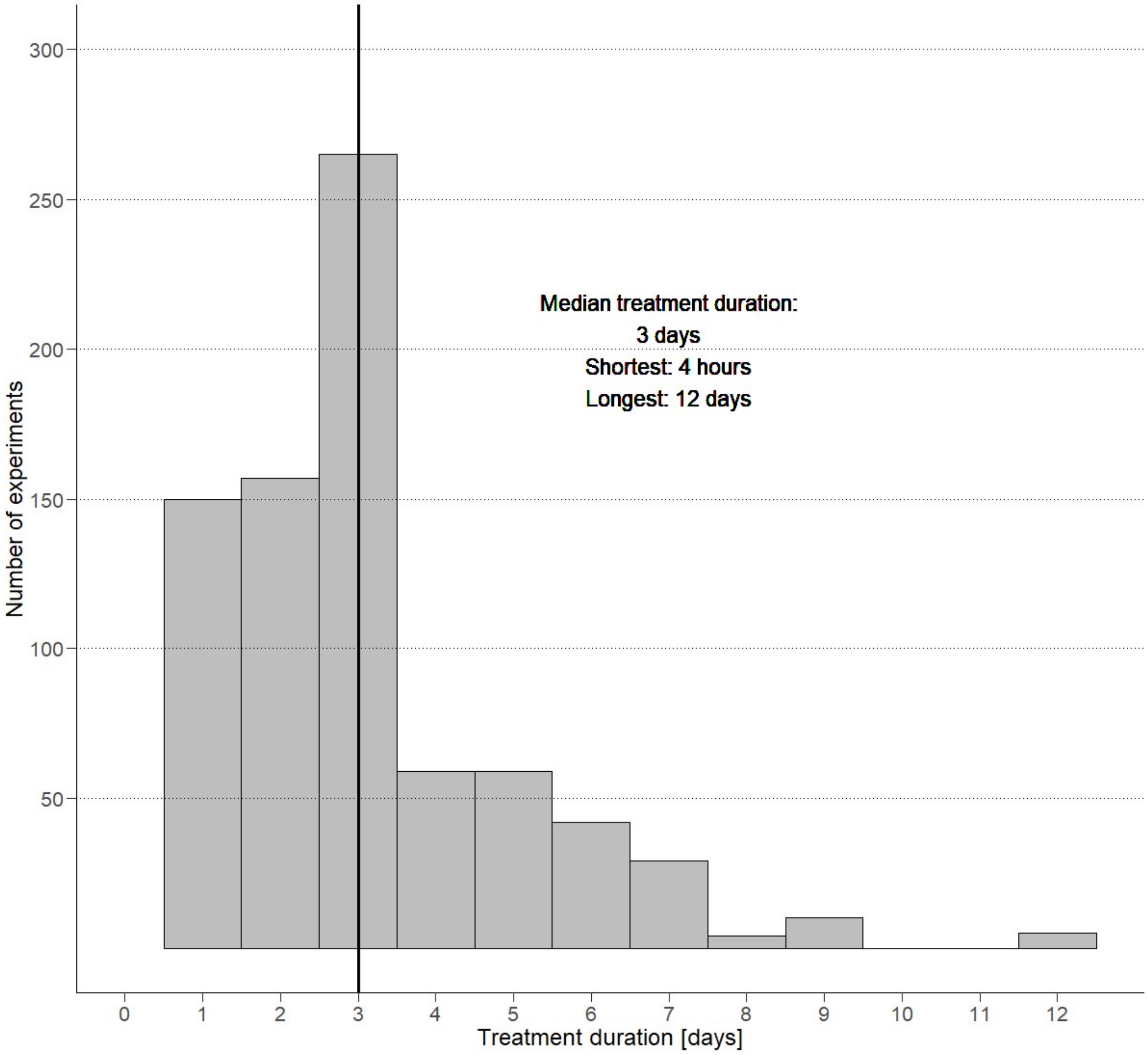


**Suppl. 8:** 137 articles included cell viability data for 786 durations of temozolomide exposure (20 unique). The lower number in comparison to used temozolomide concentrations is due to non-reporting. Treatment duration was defined as the duration of exposure of Uppsala-87 Malignant Glioma cells to temozolomide.
